# Extended perfused culture of cm-scale endocrine pancreatic tissues created through sacrificial embedded printing into alginate

**DOI:** 10.64898/2026.04.07.715214

**Authors:** Brenden N. Moeun, Hamid Ebrahimi Orimi, Théophraste Lescot, Jonathan A. Brassard, Steven Paraskevas, Sophie Lerouge, Marc-André Fortin, Richard L. Leask, Corinne A. Hoesli

## Abstract

Pluripotent stem cells represent a potentially unlimited cell source for the fabrication of human bioartificial tissues to study and treat degenerative conditions such as type 1 diabetes. Alginate is widely used for mammalian cell immobilization and the primary hydrogel studied for pancreatic islet encapsulation. Rheological properties of alginate solutions or fully gelled forms are unsuitable as support matrix for embedded 3D printing. We describe partially gelled self-healing alginate formulations tuned for embedded 3D printing. Perfusable multi-plane hierarchical networks branching into 10 parallel channels, obtained by 3D printing of Pluronic F127 into the alginate support, show high fidelity to computer-assisted models. Therapeutic β-cell doses (40×10^6^ cells/mL) within centimeter-thick perfusable constructs remained viable for at least 1 week of culture under flow, with rapid insulin secretion detected upon glucose challenges. Stem cell-derived islet clusters cultured in 5-channel contructs for 25 days differentiated towards functional insulin-expressing cells. We describe a novel approach to generate cm-scale perfusable endocrine pancreatic constructs using sacrificial embedded 3D printing into alginate. This approach offers an adaptable platform to engineer perfusable cm-scale functional endocrine pancreatic tissues and potentially other vascularized bioartificial tissues.

## Introduction

Perfused culture of human-scale (cm-thick) tissues is of substantial interest in regenerative medicine and preclinical studies – particularly with regulatory agencies shifting away from animal testing^1–3^. Achieving such tissue scales requires perfusion architectures that enable efficient mass transport. Human cells can be immobilized in hydrogel slabs, but these offer limited capacity to increase cell loading or tissue thickness without substantially increasing slab surface area^4,5^, and poorly reproduce more complex vascularized tissue architectures. Hundreds of thousands of cells can be cultured in hollow fiber bioreactors with or without hydrogel support matrices, but these do not afford control over perfusion network geometries^6,7^.

Additive manufacturing offers the opportunity to generate complex multi-material three-dimensional (3D) structures of controlled geometry. Thick tissue constructs with branched perfusion networks of controlled 3D geometry have been obtained by casting cell-laden hydrogels around self-supporting 3D printed templates^8,9^, by stereolithographic photopolymerization^7,10–13^, or by embedded 3D printing of a sacrificial material into a cell-laden hydrogel support^14–17^. Printing or support materials applied in each method are associated with different limitations. For example, self-supporting 3D printed templates such as carbohydrate glass are prone to breaking during processing or handling. Stereolithographic methods require materials that can be photopolymerized and there are concerns of damage to cells during illumination.

In embedded 3D printing, thermosensitive hydrogels such as Pluronic F127^18–20^, agarose ^21^ or gelatin ^20,22^ are printed into a self-healing cell-laden hydrogel support, after which the support is solidified and the sacrificial material is removed typically via liquefaction after a temperature shift. Successful embedded 3D printing places constraints on the support matrix, which should exhibit thixotropic behaviour to allow the matrix to deform during printing while providing sufficient mechanical support of the filament after passage of the nozzle to provide shape retention. After initial success with acellular materials^23–25^ or hydrogel baths with dilute cell doses^23,26^ this technique was more recently applied to cell spheroids to create high-density tissues that were maintained in perfused culture for up to 8 days^27^. Despite being one of the most widely used hydrogels in tissue engineering, alginate has not been considered suitable as embedded 3D matrix material since as a liquid, it is unable to support 3D printed filaments and as a gel, it is too brittle.

Alginate-based cell immobilization has been of longstanding interest in the treatment of type 1 diabetes – an autoimmune disorder targeting the insulin-producing beta cells within pancreatic islets of Langerhans^28^. In this context, alginate immobilization aims to create a size exclusion barrier between transplanted islets and the recipient’s immune system. Encapsulated islet transplantation could provide a durable alternative to daily insulin injections, while also potentially eliminating the need for lifelong immunosuppression currently applied in clinical human islet transplantation^29–31^. Encapsulation can also enable graft retrieval and localization, which is of particular interest for higher-risk cell sources of demonstrated therapeutic potential such as SC-islets)^32,33^. Alginate-based devices of different geometries such as microbeads^34–38^, fibers^39,40^ or sheets^41,42^, have been effective in protecting islet transplants from rejection for several months in diabetic animals. However, these devices have had limited success when transitioning to larger animals and thereby larger tissue constructs^18,43,44^. A major bottleneck of scaling up beta cell transplantation devices is the transport of critical molecules like oxygen and insulin^19,30^. Convection-enhanced devices anastomosed to the recipient circulation can improve transport dynamics^22,45–47^. Considering that avascular tissue constructs are restricted to approximately 200 μm in thickness^48^, the incorporation of hierarchal vascular networks can enable thicker tissues, thereby decreasing the footprint of implants and facilitating transplantation surgeries.

In this work, we present a sacrificial embedded 3D printing technique that can be used to generate thick, human-scale vascularized tissues for diabetes cell therapy using partially crosslinked alginate (**Figure 1**). We demonstrate that careful formulation leads to alginate matrices with thixotropic behavior suitable for embedded 3D printing to create complex hierarchical multi-plane channel networks. Dispersed beta cells, beta cell clusters, or SC-islets can be immobilized at high densities and maintained viable for weeks under perfusion, with efficient insulin response kinetics upon glucose challenge.

**Figure 1.**
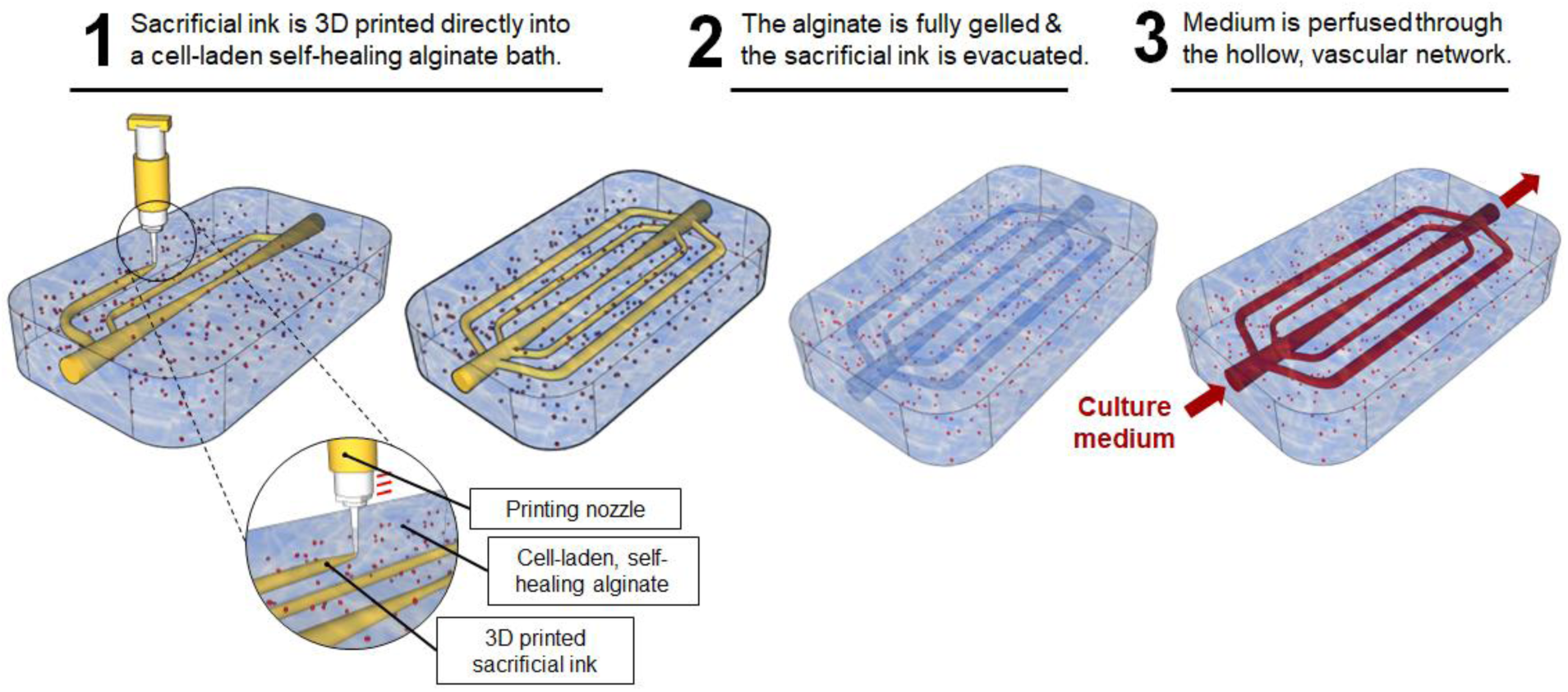
Vascularization of thick beta cell-laden tissue constructs via embedded 3D printing into a cell-laden, self-healing alginate. First, a vascular template is 3D printed directly into the partially gelled alginate matrix. The alginate can then be fully gelled using additional crosslinker to immobilize the 3D printed materials and cells. Next, the sacrificial vascular template is evacuated to leave behind hollow channels that can be perfused with cell medium or blood.

## Results

### Self-healing alginate formulation

An embedded 3D printing matrix requires a thixotropic property whereby the material behaves like a solid at zero to low shear but like a liquid with enough exerted shear stress. The threshold between these two behaviors is the yield stress (τ_0_). Conventionally, alginate can exhibit entirely liquid behavior (i.e., with no crosslinker) or entirely solid behavior (i.e., with crosslinker). By gradually gelling alginate solution with limiting concentrations of calcium ion crosslinker (10.0-20.0 mM) under constant agitation, we were able to formulate thixotropic formulations (**Figure 2A**). By fitting a Herschel-Bulkley model on the rotational rheology data, we empirically determine the yield stress, flow coefficient (k), and Herschel-Bulkley index (n) of the alginate formulations (Supplementary **Table S1**). All formulations demonstrated shear-thinning behavior (n<1). Using more than 20.0 mM of calcium for partial gelation led to an increased frequency of heterogeneous gelation likely due to heightened gelling kinetics. The yield stress was also estimated from the flow point (intersction between the storage and loss modulus) in oscillatory rheometry^49^ (**Figure 2B**).

**Figure 2.**
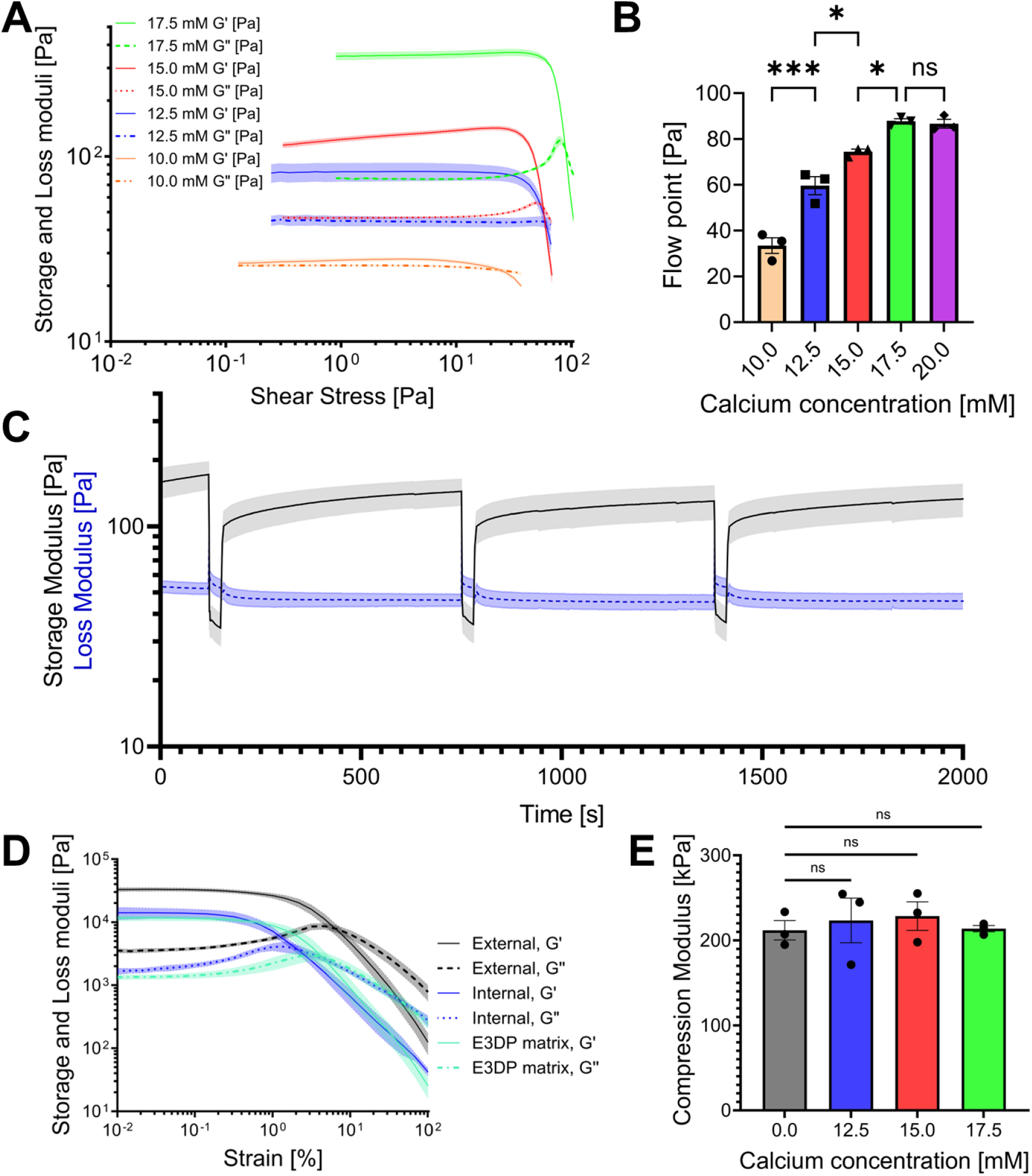
Rheological and mechanical properties of self-healing alginate matrices. **(A)** Storage (solid lines) and loss (dotted and dashed lines) moduli of alginate partially gelled with 10.0-17.5 mM of calcium. **(B)** Flow points of partially gelled alginate. **(C)** Recovery of alginate partially gelled with 15.0 mM of calcium after step increases of shear stress beyond the flow point. Solid line represents the storage modulus, and the dotted line represents the loss modulus. **(D)** Shear moduli of alginate formulations after complete gelation. G’: storage modulue. G’’: loss modulus. **(E)** Compressive moduli of alginate formulations after complete gelation. Error bars and shaded areas represent the standard error of the mean and statistical significances is indicated as by *P<0.05, **P<0.01, and ***P<0.001.

As more crosslinker is added, there is a general increase in plateau storage modulus, plateau loss modulus (**Figure 2A**), and yield stress/flow point (Supplementary **Table S1**, **Figure 2B**), confirming that rheological properties are dependant on the calcium concentration. Considering that the yield stress of matrix materials for embedded 3D printing should be at least one order of magnitude less than that of the 3D printed inks^50^, these results indicate that partially gelled alginate can be tuned for a variety of sacrificial inks.

To evaluate whether these formulations could support the embedded 3D printing of complex vascular networks, we performed recovery tests. In this analysis, partially gelled alginate formulations were subjected to step increases in shear stress and returned back to a low shear state. This stepwise increase/decrease was repeated multiple times to simulate a translating printing needle passing through the same location when building something complex like an omnidirectional vascular network. **Figure 2C** shows that alginate partially gelled with 15.0 mM of calcium demonstrates an almost immediate transition from a solid to liquid state in response to a step increase in shear stress, and instantly reverts back to a solid-like state when shear is decreased. However, the recovery of the storage modulus after yielding required several minutes (**Figure 2C**).

Importantly, the alginate matrix must be capable of fully solidifying after embedding the vascular template, thereby enabling efficient removal of the sacrificial ink and formation of a hollow perfusion network (**Figure 1**). After complete gelation, our alginate formulations exhibit an increase in plateau storage and loss moduli by around 2 orders of magnitude. These values are comparable to the moduli of internally gelled alginate without a prior partial gelation step (**Figure 2D**). When comparing the compression modulus of our alginate formulations after complete gelation with alginate that was not previously partially gelled, we measure no statistical difference (**Figure 2E**) suggesting that our perfusable tissue will be stiff enough to withstand the forces associated with pressure-driven flow during perfusion.

### Sacrificial embedded 3D printing into alginate

Having identified promising self-healing alginate candidates that could be used for embedded 3D printing, we evaluated their capacity to generate perfusable constructs. Although multiple formulations showed thixotropic behavior, alginate crosslinked with less calcium demonstrated a yield stress too low to keep filaments in place throughout the embedded 3D printing process (**Figure 3A** and **B**). Meanwhile alginate that was overly stiff (e.g., crosslinked with 20.0 mM calcium or more) demonstrated a greater tendency to result in nonhomogeneous gelling and constrict printed filaments, ultimately decreasing print fidelity. Formulations crosslinked with 15.0-17.5 mM calcium supported the printing of multibranched vascular template with a variety of filament angles. As expected from mass balances around the nozzle, the hydraulic diameter of the resulting channels was proportional to the square root of the ratio between the extrusion speed and the nozzle displacement speed. With 20G diameter print nozzle used in our studies, the channel hydraulic diameter was tuned between ∼0.5 mm and 2 mm simply by adjusting the nozzle displacement speed and extrusion rate (**Figure 3C**). This level of control is critical for tuning hydraulic resistance and maintaining balanced flow distributions across complex branching networks.. The higher density of calcium-containing alginate gels compared to water-filled channels created strong contrast during μCT scanning, enabling clear visualization of both components. This contrast enhancement, provided by the calcium content in the alginate, allowed precise delineation of the interface between the alginate matrix and the channels (**Figure 3D**; see methodology for details). The resulting contours of the alginate construct after vascular 3D printing confirmed that the self-healing alginate formulations successfully supported the fabrication of vascularized constructs. As shown in the reconstructed images from segmented μCT images (**Figure 3D**), the branching channels were well connected. These findings demonstrate the ability to control channel diameters, achieve high print fidelity, and maintain structural integrity without cracks larger than 45 μm around the channels (**Figure 3E-I**).

**Figure 3.**
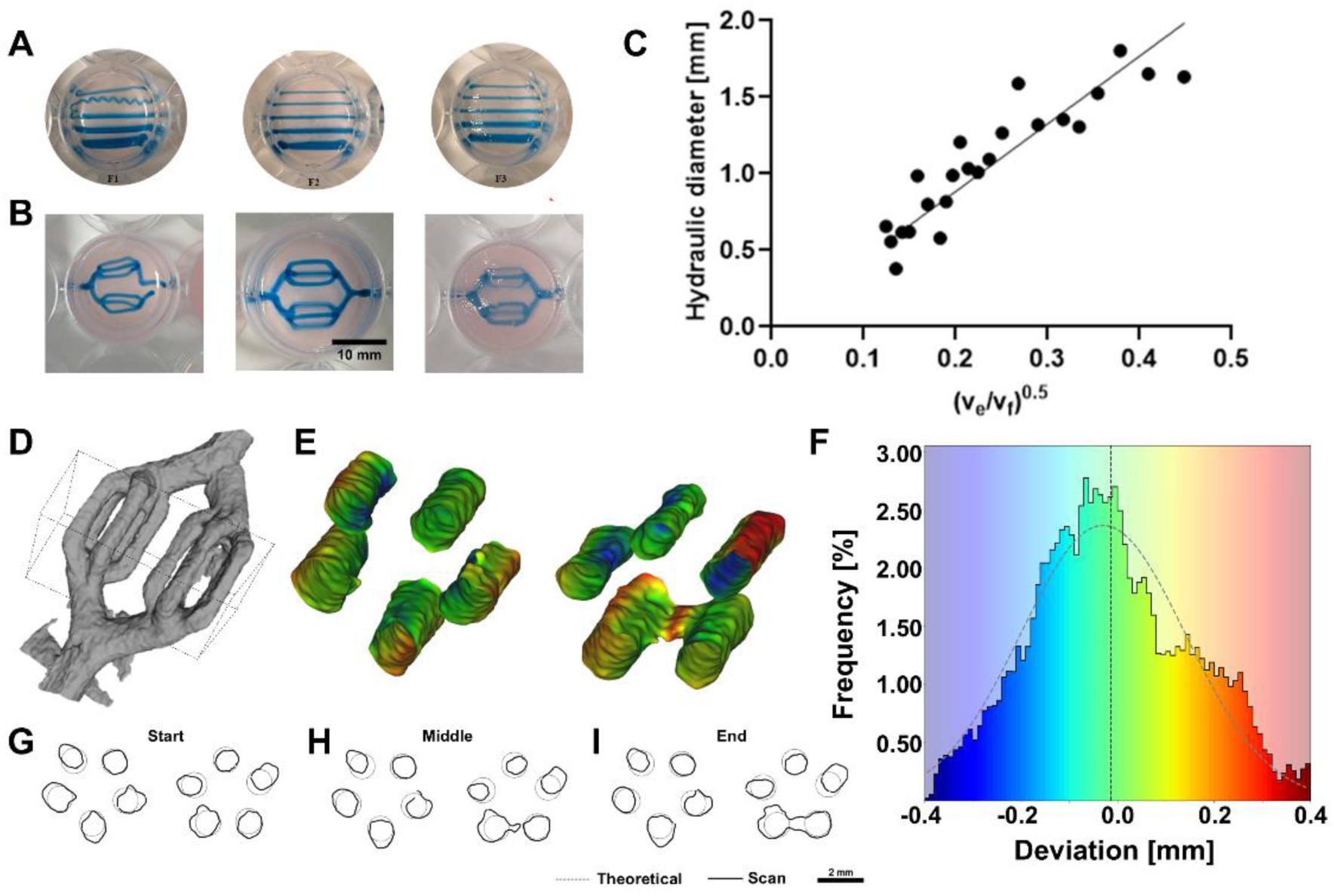
Evaluating the embedded 3D printing of vascular templates and perfusable networks within self-healing alginate formulations. **(A-B)** Embedded 3D printing of 35% Pluronic F127 coloured with blue dextran into alginate partially gelled with 10.0 mM, 15.0 mM, and 20.0 mM of calcium. **(A)** Modulating print speed to vary the diameter of 3D printed filaments (going downwards: 3.00 mm/s, 2.00 mm/s, 1.00 mm/s, 0.75 mm/s, 0.50 mm/s). **(B)** 3D printing sacrificial vascular templates with branching geometries. **(C)** Hydraulic diameters of channels as a function of print speed (v_p_) and extrusion rate (v_e_). **(D)** 3D model segmented from µCT scans of a 10-branch perfusable network in alginate. **(E)** Heat map showing minor (green, 0–0.10 mm), intermediate (yellow and cyan, 0.10–0.25 mm), and major (red and blue, ≥0.25 mm) deviations of the 10 branching channels when compared with the intended geometry. **(F)** Histogram of deviations throughout the comparative analysis, normalized to frequency (mean = -0.014 mm, σ = 0.16 mm, black dashed line; Gaussian mean = -0.026 mm, σ = 0.16 mm, grey dashed line). **(G-I)** Cross sectional comparisons of the 10 branching channels at the **(G)** start, **(H)** middle, and **(I)** end.

### Cell viability and function

To assess cell survival during biofabrication, we evaluated the impact of the embedded 3D printing process on mouse insulinoma 6 (MIN6) beta cell survival at different stages of the embedded 3D printing process. **Figure 4A** shows that the majority of MIN6 cells remain viable after the mixing (∼1 min), embedded 3D printing (<5 min), complete gelation (∼15 min), chilling (∼10 min) , and washing steps (∼1 min) of the sacrificial embedded 3D printing process. There were no changes in viability when comparing free MIN6 cells with those immobilized within unmodified alginate and those immobilized within previously partially gelled alginate. MIN6 viability was also unaffected by the embedded 3D printing process with Pluronic F127. Using an internal gelation final solidification step with additional calcium carbonate and GDL also did not influence MIN6 viability when compared to using an external gelation step with a calcium chloride solution (**Figure 4B**). Similarly, stage 3 posterior foregut cell clusters showed high levels of viability after the biofabrication steps (**Figure 4C**).

**Figure 4.**
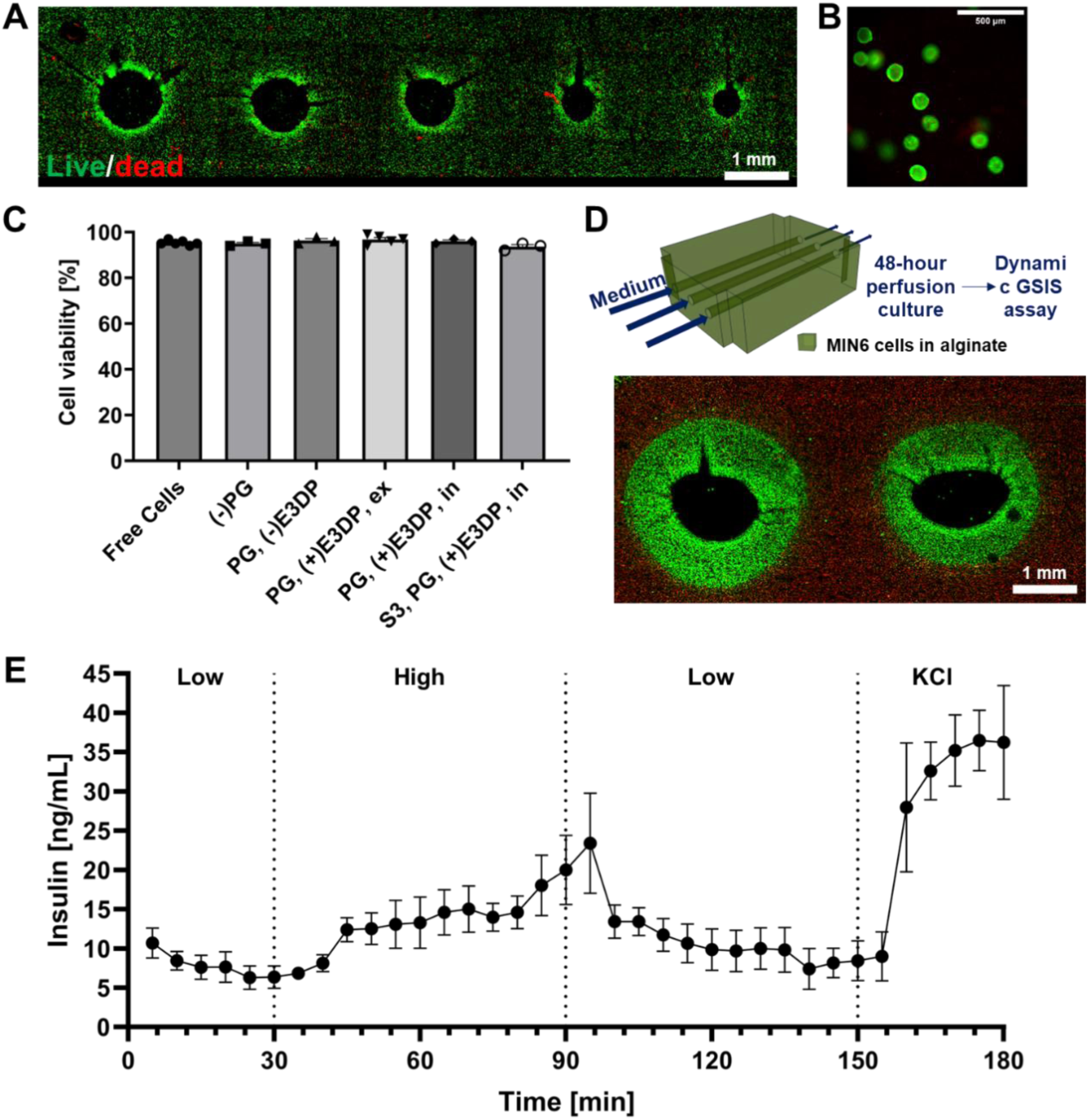
Viability and function of pancreatic cells within perfusable alginate constructs generated via sacrificial embedded 3D printing within self-healing alginate. **(A)** Live (green, calcein-AM) and dead (red, propidium iodide) staining of MIN6 single cells (40·10^6^ cells/mL) after the sacrificial embedded 3D printing process. Channel diameters are controlled by varying print speed (from left to right: 1.0 mm/s, 1.5 mm/s, 2.5 mm/s, 3.5 mm/s, 4.5 mm/s). **(B)** Viability stain of S3 posterior foregut clusters after sacrificial embedded 3D printing process. **(C)** Quantification of cell viability after the sacrificial embedded 3D printing process. (PG: partial gelation, E3DP: embedded 3D printing with Pluronic F127, ex: external gelation final step with CaCl_2_, in: internal gelation final step with CaCO_3_, S3: dissociated stage 3 posterior foregut clusters derived from PSCs). **(D)** Top: schematic of experimental setup. Perfusable beta cell constructs with 3 straight channels are cultured under flow for 2 days and subsequently subjected to a dynamic GSIS assay. Bottom: beta cell viability around 2 of the channels within the constructs (starting density: 40·10^6^ MIN6 cells/mL) after dynamic GSIS testing. **(E)** Dynamic GSIS function of perfusable constructs after 2 days of perfusion culture (3.3 mL/min/channel).

We compared MIN6 survival profiles under perfusion to our previous observations in static immobilized culture^4^ as well as simplified perfused culture models^9^. After 2 days of perfusion, only cells within 600 ± 85 μm of a perfusable channel remained viable (**Figure 4D**). Cells further away lack an adequate source of oxygen. This viable region coincides with what we previously have reported for the same cell seeding and culturing conditions^9^. The limited region in which cells can remain viable highlights the need for more complex vascularization and a denser perfusable network.Insulin secretory response dynamics are a key consideration towards applications in diabetes research or cellular therapy.

Glucose-stimulated insulin secretion (GSIS) of beta cell-laden alginate constructs was evaluated after 2 days of perfusion culture. Following a step increase in glucose, we measured a rapid insulin release response that reverted to pre-stimulation levels in response to a return to the basal glucose condition (**Figure 4E**). This dynamic response is consistent with what others have reported with macro- and microencapsulated beta cell lines under perfusion^51,52^. Depolarization with KCl also led to a spike in insulin secretion. Of note, a ∼10-min delay in response to each step change in glucose concentration was observed. Considering a dead volume of ∼50 mL within the flow loop and a 10 mL/min flowrate approximately 5 min of this delay can be attributed to fluid residence time of the perfusion system.

### Constructs with branching vascular networks

We expanded on the technology’s capabilities by using more complex vascular geometries to maintain the viability of therapeutic cell densities, prolonging culture times, and employing more physiologically relevant cell models. We started by loading our 10-branch tissues **Figure 4A-B**) with a concentration of MIN6 cells (40 × 10^6^ cells/mL) of oxygen consumption rates equivalent to 50% of a therapeutic adult human islet dose (moderate metabolic demand model; Supplementary **Table S2**). **Figure 5C-F** demonstrate that, throughout the cm-scale constructs, only cells around the channel lumens remained viable after 2 days of perfusion culture. Comparing these viable regions with our computational model, we obtain moderate agreement, with some viable regions spanning beyond the expected theoretical oxygen tension threshold of 1.16 mmHg which was identified in our previous work^9^ (**Figure 5C**). This could be the result of deviations in vascular network geometry—namely, variations in channel diameter and spatial coordinates from the intended specifications—as well as alginate swelling, which can further alter channel dimensions and actual cell concentration, and the exclusion of possible axial oxygen transfer within our 2D mass transport model.. Excluding the viability regions arising from superficial oxygen diffusion, beta cell viability is sustained for an additional ∼5 mm of depth within these vascularized constructs.

**Figure 5.**
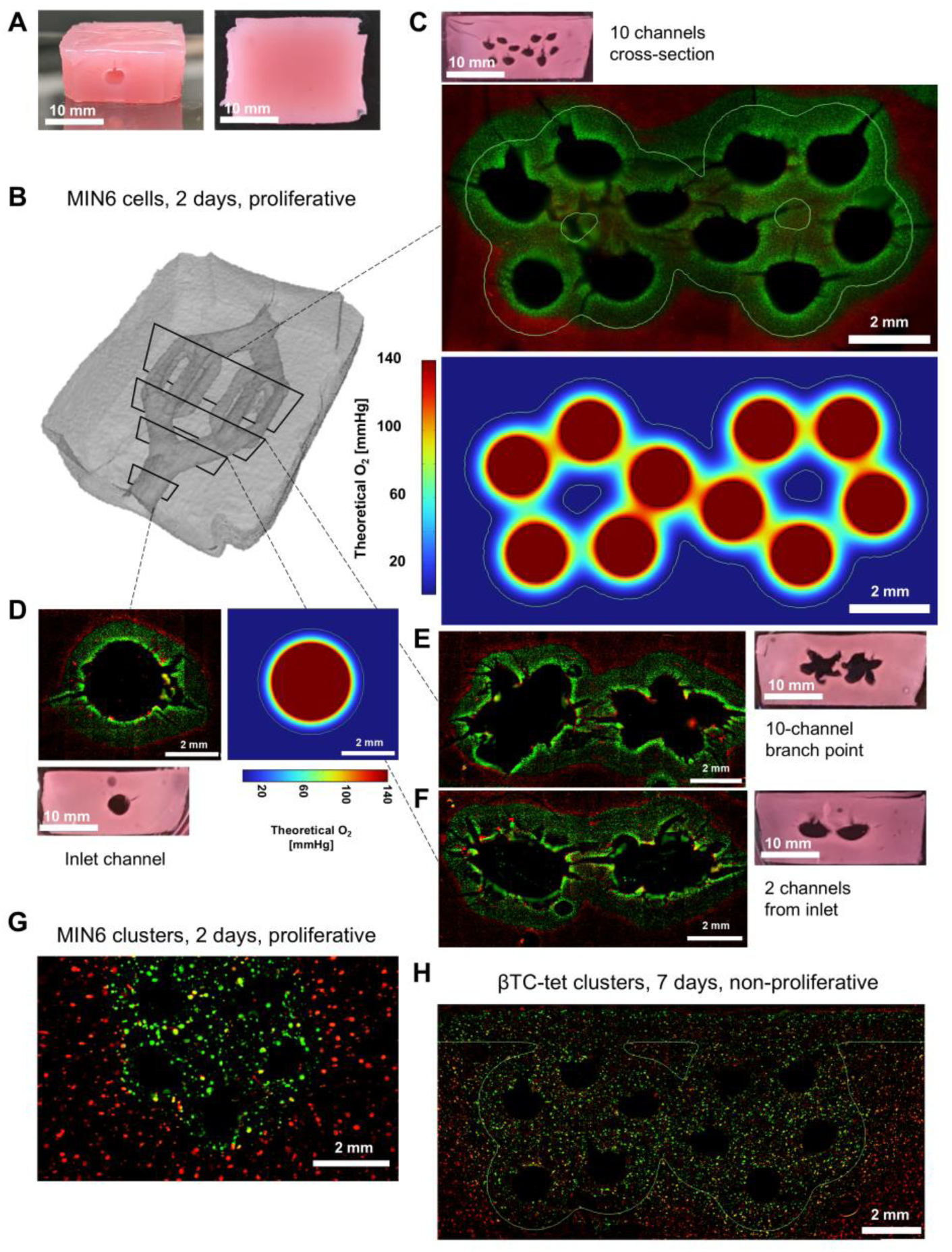
Days- to week-long culture of cm-scale beta cell-laden constructs with multi-branched vascular networks under flow. **(A)** Whole constructs after perfusion culture and before sectioning. Scale bars are 1 cm. **(B)** 3D rendering of an alginate construct irrigated with a 10-branch vascular network indicating the approximate location of subsequent cross-sectional slices used for viability staining. **(C-F)** Live (green, calcein-AM) and dead (red, propidium iodide) staining of 10-branch construct sections after 2 days of perfusion culture (starting cell density: 40·10^6^ single MIN6 cells/mL). **(C)** Left: cell viability around 10 channels with theoretical viability threshold contour (1.16 mmHg, green line. Right: corresponding computational oxygen transport model. **(D)** Viability of cells around the branch point where 2 channels divide into 10. **(E)** Cell viability around 2 channels originating from inlet. **(F)** Left: cell viability around inlet channel. Right: corresponding computational oxygen transport model. **(G)** Viability of MIN6 clusters (starting cell seeding: 4000 clusters/mL, 500 cells/cluster) in a 5-branch construct after 2 days of perfusion culture with theoretical viability threshold contour (1.16 mmHg, green line) **(H)** Top: viability of βTC-tet clusters (starting cell seeding: 31,000 clusters/mL, 1100 cells/cluster) within a branching 10-channel construct after 7 days of perfusion culture with theoretical viability threshold contour (1.16 mmHg, green line).

Moving towards a more physiologically relevant comparison to islets, which are natively cell clusters, we immobilized MIN6 clusters within our perfusable tissues. After 2 days of perfusion culture, we observed an expected viability pattern where cell clusters around the 5-channel perfusion network are well oxygenated and viable while clusters far away, do not survive (**Figure 5G**). Regarding culturing times and oxygen consumption estimations, the use of MIN6 cells as models for insulin-producing beta cells can be limiting due to their relatively rapid doubling times^51^ when compared to primary beta cells which, in humans, are long-lived^53^. In addition, cell-specific growth rate correlates positively with oxygen consumption rate, as proliferating cells exhibit higher metabolic and mitochondrial activity than quiescent cells^54,55^. Recognizing that primary beta cells have a very low turnover rate^56^ we pursued a more physiologically relevant cell model by using a therapeutic concentration of βTC-tet clusters under growth arrest. This established a more reliable steady state, increase culturing times and minimize the changes in oxygen consumption dynamics caused by cell growth. After 7 days of perfusion culture, the regions of high cell viability followed an expected pattern of localizing around the perfusion channels **Figure 5H**). Further, these results align with our computational model, with most viable cell clusters falling within the outlined theoretical viability limit.

### Vascularized stem cell-derived pancreatic tissue constructs

After establishing a robust biomanufacturing and culturing platform using cell lines, we evaluated whether SC-derived cell clusters can be maintained viable under long-term perfusion in cm-scale constructs without compromising SC-islet maturation. Immature SC-islets were obtained through directed differentiation with transition from adherent culture (Stages 1-4) to microwell-based aggregation (end of Stage 4), microwell culture (Stage 5) and suspension culture (Stage 6) (**Figure 6A**; Supplementary **Figure S1**)^57^. During the differentiation, the cells progressed through the expected endoderm (>95% SOX17^+^) and pancreatic progenitor (>75% PDX1^+^/NKX6.1^+^) intermediates (**Figure 6A** ii). Controls maintained in suspension for a further 25 days (Stage 7 day 27) give rise to endocrine clusters (**Figure 6A** iii). As expected, these clusters can be stained with dithizone, a chelator of Zn^2+^ found in insulin granules (**Figure 6A** iv).

**Figure 6.**
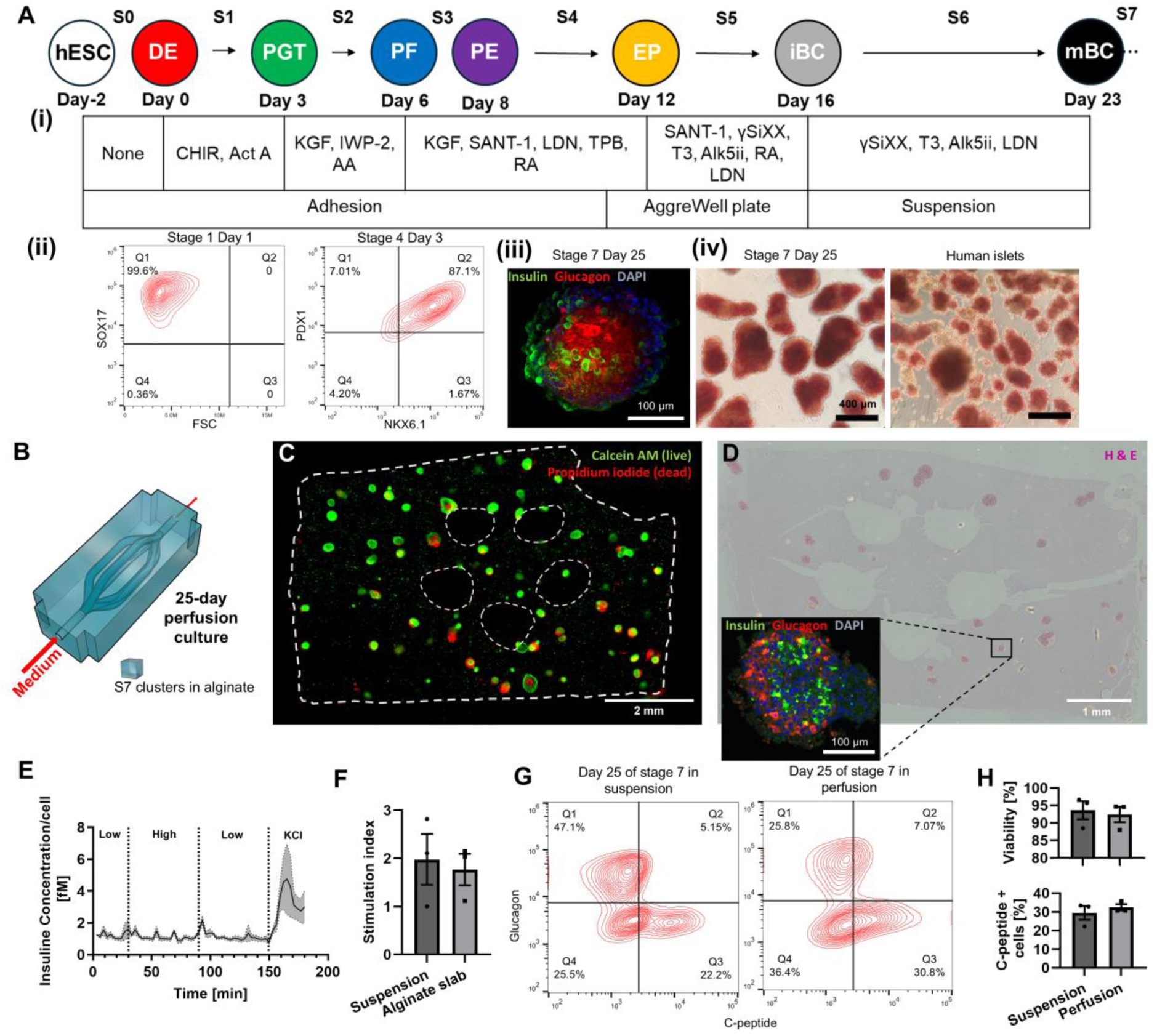
Long-term perfused culture of stem cell–derived endocrine pancreatic tissues. This figure illustrates the generation, fabrication, and functional assessment of vascularized pancreatic constructs derived from human pluripotent stem cells. **(A)** A schematic that outlines the stepwise differentiation protocol leading to insulin-producing β-like cells (i), with sample flow cytometry profiles indicated for Stage 1 and Stage 4 (ii). Representative immunofluorescence images of Stage 7 clusters (iii) along with dithizone-stained stage 7 cluster and human islets (iv) are shown to highlight endocrine cell morphology. **(B)** 3D rendering of tissue construct with hierarchical perfusable channels **(C)** Live/dead staining of a cross-section from the printed construct after 25 days of perfusion culture in Stage 7. **(D)** Histological analysis following 25 days of perfusion includes hematoxylin and eosin (H&E) staining and immunofluorescence for insulin, glucagon, and DAPI. **(E)** Dynamic glucose-stimulated insulin secretion (GSIS) of the perfused constructs after 25 days with **(F)** static controls. **(G)** C-peptide and glucagon co-expression in Stage 7 cells of perfused constructs (right) and suspension (left) after 25 days of culture. **(H)** Quantitative analysis of C-peptide–positive cells and cell viability in perfused constructs compared to suspension controls.

To study SC-islet maturation under perfusion, Stage 6 SC-islet clusters were immobilized within a 6-channel alginate constructs (**Figure 6B**). After extended perfusion culture (Stage 7 day 25), the cell clusters remained highly viable and the overall structural integrity of the cell-laden hydrogel and its channels was preserved (**Figure 6C**). The 25-day Stage 7 alginate-immobilized culture period led to SC-islet maturation, as shown by numerous monohormal insulin-positive (β-like) cells and glucagon-positive (α-like) cells present within cell clusters throughout the constructs (**Figure 6D**) – analogous in compposition to the suspension controls (**Figure 6A** iii). Stage 7 perfused tissue function was assessed via GSIS (**Figure 6E**). While glucose stimulation did not result in as clear of an insulin secretory response at the device outlet as compared to MIN6 cells, depolarization with KCl induced clear insulin release. The delay in response to glucose stimulation was also observed in primary human islets (Supplementary **Figure S3**), suggesting that the encapsulation environment rather than the intrinsic cell maturity explains the kinetics. Furthermore, SC-islets recovered from perfusion constructs showed maintained glucose responsiveness in static incubation assays (**Figure 6F**).

When comparing SC-islets recovered from the perfused constructs to suspension controls, no statistically significant differences were identified with respect to cell viability nor the proportion of C-peptide^+^ (32 ± 2% in perfusion vs 29 ± 5% in suspension) or glucagon^+^ (23 ± 2% vs 38 ± 9%) cells by flow cytometry (**Figure 6G-H**). In both culture systems, we did not identify substantial co-expression of these markers, contrary to other groups who have reported bi-hormonal populations which may represent early endocrine populations that do not substantiall contribute to adult endocrine cells in normal human development^59–61^.

## Discussion

The sacrificial embedded 3D printing approach developed in this work offers a simple and clinically aligned way to build thick, perfusable tissue constructs using materials adaptable to clinical-grade production. Alginate is already deeply established in translational cell therapy because of its safety, ease of processing in physiological conditions, low immunogenicity and potential for creating immunoprotective size exclusion barriers^62^. However, in its native form alginate does not flow or self-heal in the way that embedded printing requires. By partially crosslinking the polymer, we created thixotropic, self-healing alginate formulations that behave like a liquid during printing but regain their structure immediately afterward. This property eliminates the need for complex composite support materials, many of which are difficult to translate clinically. With this approach, we reliably produced cm-scale constructs containing up to ten perfusion channels (**Figure 4** and **Figure 5**). Alginate construct and channel integrity was maintained during long-term culture delivering oxygen and nutrient to immobilized cells. The ability to combine sacrificial embedded printing with a clinically accepted hydrogel represents a step forward in the practical manufacturing of vascularized artificial tissues, particularly for applications where thickness and high cell density are essential.

The partially gelled alginate used here also worked as an extrusion bioink, producing multilayer structures and supporting viable beta cells after printing (Supplementary **Figure S4**). This dual functionality is useful for workflows where sacrificial printing and direct cell printing need to be combined. Similar advantages of partially gelled alginate have recently been reported elsewhere for fibroblast-laden constructs^63^. Our systematic method for tuning gelation could likely be extended to other hydrogels such as collagen, which remains one of the most used biomaterials in tissue engineering but is notoriously difficult to print without chemical or mechanical modification^64, 65, 66^. Controlled partial gelation strategies, including those using tannic acid^67^, may allow the embedded-printing concept to expand beyond alginate toward other ECM-based systems.

The perfusion networks generated using the partial gelation and embedded 3D printing method reproduced branching architectures that mimic essential features of vasculature (**Figure 3**). While print fidelity was generally high, some inconsistencies were observed, especially in small-diameter channels where the printing needle moves more quickly. These included teardrop-shaped cross sections, occasional channel fusion, and spatial drift (**Figure 3G-I**). Many of these issues are known in sacrificial embedded printing and result from the forces exerted by the moving print head^27,68^ or from changes in the gel during final crosslinking and perfusion^69, 70^. These imperfections did not prevent long-duration perfusion or compromise cell viability. Future optimization of print speed, extrusion rate, and matrix rheology could help achieve more consistent geometries. Despite these engineering challenges, the constructs shown here represent, to our knowledge, the first example of perfusable, branching vascular networks supporting cm-thick β-cell-laden tissues for multiple weeks.

Maintaining oxygenation is a central challenge when engineering thick tissues, particularly when using cells with high oxygen consumption rates such as pancreatic islets^71^. Therapeutic doses for type 1 diabetes require hundreds of thousands of islet equivalents—roughly 0.5-1 billion cells^72,73^—and delivering this number of cells within a compact device requires high seeding densities. In traditional encapsulation systems, diffusion alone cannot meet the oxygen demand of such densely packed, metabolically active cells. By contrast, our constructs, seeded at 40×10⁶ MIN6 beta cells/mL, approached clinically relevant volumes and volumetric oxygen consumption rates, and yet maintained viability across nearly 1-cm depths for at least 7 days in standard cell culture conditions (5% CO_2_, ∼140 mmHg pO ^74,75^ **Figure 5**). This demonstrates that perfusable channels can help overcome the mass-transport limitations that have historically restricted the thickness of engineered endocrine tissues. If the oxygen concentration in the perfused fluid is lower – as would be the case in physiological environments – higher channel densities or lower cell densities would be required to maintain full tissue viability^4,9^. Higher channel densities could be achieved via further printing process optimization, or by introducing multi-step 3D printing approaches that incorporate angiogenic materials and secondary channels within the alginate.

Functionally, the perfused 3-channel constructs with single MIN6 cells exhibited rapid and reversible response to a step change in glucose concentration (**Figure 4**). The ∼5-min response time meets the <15-min requirement suggested for maintaining safe postprandial glucose control^76^ and closely matches the kinetics of primary human islets cultured in vitro^77^. The performance is also comparable to intervascular devices tested in vivo^78^ and approaches the onset times of fast-acting insulin analogs^79^. Increasing the density or complexity of the channel networks—such as those shown in **Figure 5** and **Figure 6**—could likely improve these kinetics even further. These results suggest that full vascularization of each islet is not strictly necessary; rather, well-distributed perfusion channels can enable fast glucose sensing and insulin release across large volumes of tissue.

A major goal of this study was to determine whether this platform could support SC-islets, a leading source for future diabetes cell therapies. Extended perfused culture of immature SC-islets in 5-channel cm-scale alginate constructs led to high survival and successful maturation into monohormonal C-peptide^+^ (co-secreted with insulin) and glucagon^+^ cells. While blunted or highly delayed response to glucose stimulation was observed *in situ* (**Figure 6E**), SC-islets recovered from constructs were glucose responsive (**Figure 6E**). It is interesting to compare insulin response dynamics between SC-islets seeded at lower densities (∼5000 aggregates/mL alginate ≈ 7.5 × 10⁶ cells/mL cells/mL, **Figure 6**) with MIN6 cells at higher seeding densities (40 × 10^6^ cells/mL). The average glucose and insulin diffusion distances to each beta cell would be substantially lower with single MIN6 cells which are in higher numbers in the channel vicinity and which are more uniformly distributed throughout the hydrogel. Beyond diffusion limitations and oxygen consumption attributed to fibrotic overgrowth on macroencapsulation devices^44,80^, our observations suggest that other factors such as beta cell distribution within devices or even islet diameter distributions should also be considered. The embedded 3D printing approach could help explore the effect of different vascular densities and cellular distributions on insulin response dynamics experimentally. In a pilot study performed with earlier (Stage 3) cells, similar viability profiles were observed (Supplementary **Figure S2**) suggesting that full in situ differentiation at earlier stages may also be feasible. Other interesting applications of our platform would be to utilize the system for process intensification, whereby large numbers of SC-islets could be cultured long-term in highly compact tissue cultures. Different alginate concentrations, modified alginates or blends with other biopolymers could be explored to drive further SC-islet maturation and function, or to introduce channel endothelialization to explore endothelial-endocrine interactions under different channel densities.

In summary, this work presents a clinically aligned, scalable, and versatile strategy for fabricating thick, perfusable tissues using sacrificial embedded 3D printing in GMP-compatible alginate. This approach can support high-density SC-derived islet tissues with long-term viability, preserved endocrine identity, and sustained functional potential. The constructs avoided the central necrosis commonly seen in macroencapsulated tissues and supported both early-stage progenitors and late-stage endocrine clusters. While further improvement of response kinetics is possible, the overall performance demonstrates how integrating perfusable channels into GMP-compatible hydrogels can address major challenges in β-cell replacement therapy. Beyond β-cell applications, the approach has broad utility. Alginate is widely used to engineer liver, cardiac, skin, and other tissues, and many of these systems struggle to maintain viability beyond a few hundred micrometers. Embedded perfusion networks provide a path toward thicker, more physiologically relevant engineered tissues ⁷⁷. The platform could also support advanced organoid culture, 3D cancer models, and drug-testing systems that require sustained nutrient exchange.

## Methods

### Alginate partial gelation

Alginate formulations were prepared from an autoclaved 6% w/v stock solution of alginate (IFF Nutrition & Biosciences through DuPont, Protanal LF 10/60) in HEPES buffered saline (pH 7.4, 10 mM HEPES, Fisher Scientific, BP310 and 170 mM sodium chloride, NaCl, Fisher Scientific, BP3581). In a 150-mL beaker, the alginate stock was mixed with additional filter-sterilized HEPES buffered saline and autoclaved 0.5 M calcium carbonate suspension (CaCO_3_, VWR, 130101) to reach a final CaCO_3_ concentration of 45 mM. The solution was kept under constant agitation (600 RPM) on a magnetic stir plate using a 1.5-inch magnetic stir bar. A 0.48 M glucono-δ-lactone (GDL, MilliporeSigma, G4750) solution was quickly prepared by vortexing the GDL in HEPES buffered saline. The GDL solution was immediately filter sterilized and added into the stirring alginate solution according to the desired degree of partial gelation in a 2:1 ratio of moles of GDL to moles of CaCO_3_. The mixture was covered and left to gel at room temperature overnight under constant stirring (100 RPM) in a biosafety cabinet. After all additions, the final alginate concentration was 3.33 % w/v.

### Rheometry of partially gelled alginate

Prior to testing, 10% v/v of cell medium was mixed into the partially gelled alginate, yielding a final alginate concentration of 3% w/v. Acellular alginate formulations were studied using a modular compact rheometer (MCR 302, Anton Paar) equipped with a cone and plate geometry (25 mm diameter, angle β=1°, CP25-1). A 0.049-mm gap was set at the truncated tip and testing was conducted at ambient conditions (∼22°C). Shear stress (𝜏) as a function of shear rate (𝛾̇, 0.01 s^-1^ to 125 s^-1^) was measured and the Hershel Bulkley model (**Equation 1**) was fitted onto these data to approximate the yield stress (𝜏_𝑜_), flow coefficient (𝑘), and Hershel Bulkley index (𝑛). Thixotropic properties were measured using oscillatory tests performed at a frequency of 1 Hz (0.1% to 120% strain). Flow points were measured at the intersection of the storage and loss moduli curves.

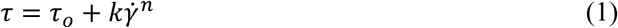

Thixotropic recovery was evaluated using a sequence of oscillatory tests. First a 1% strain was exerted on the material for 2 min, then a shear stress step increase to 110% of the flow point was applied for 0.5 min. Next, the applied stress was returned to 1% strain for 10 min. This was repeated two additional times per sample.

### Mechanical properties

As a final gelation step, a concentrated GDL solution (2.25 M) was added to react with the remaining CaCO_3_. Gels were then quickly casted between 2 glass slides separated by spacers (1 mm for shear moduli; 2 mm for compressive moduli) and left to gel for 20 min. Afterwards, gels were placed in 100 mM calcium chloride (CaCl_2_, MilliporeSigma, 1023780500) solution for 10 min and transferred into cell culture medium supplemented with an additional 5 mM of CaCl2 overnight to swell. The gels were then cut into 25-mm disks with a circular cutter. Oscillatory frequency sweeps with a sandblasted 25-mm parallel plate geometry (Anton Paar, 3997) were used to measure the shear moduli of the fully gelled alginate. The hydrogels were compressed by axial force (∼0.2 N) and the applied oscillation strain ranged from 0.01 to 100% using an angular frequency of 10 rad/s. Stress-strain plots were also generated from the unconfined compressions (5 μm/s) of fully gelled discs. Strain was defined as 1-H/H0 where H is the height of the top plate and H0 is the initial height. The normal force was divided by the plate area to calculate the stress and linear regression was used to measure the compressive modulus of the gels.

### Sacrificial ink preparation

Pluronic F127 (MilliporeSigma, P2443) powder was mixed with distilled water (Thermo Fisher Scientific, 10977015) to yield a final concentration of 35% w/v. This mixture was left in a 4°C fridge overnight to dissolve. For cell experiments, dissolved Pluronic F127 ink was autoclaved for 30 min. When evaluating suitable matrix materials, the Pluronic F127 ink was formulated with 2.9 mg/mL blue dextran (2000 kDa, MilliporeSigma, D5751) prior to printing to visualize the printed filaments.

### Capacity for embedded 3D printing

To test for suitable embedded 3D printing matrices, 1 g of acellular alginate formulations, prepared as described above were scooped into the middle wells of an untreated 12-well plate (Sarsedt, 833921500) using a metal spatula. The plates were then centrifuged for 2 min at 300 x g to remove air bubbles. Afterwards, blue sacrificial ink was used to 3D print filaments at varying translational speeds and branching networks directly into the alginate (3D Discovery, RegenHU). The capacity to support these structures was evaluated qualitatively.

To characterize the capacity to form hollow perfusable networks, an additional GDL solution (0.4 g/mL) was first mixed into the alginate formulations to react with the undissociated CaCO_3_ (2:1 molar ratio) which was calculated to be 45 mM minus the amount that was dissociated for partial gelation. The alginate was then quickly scooped into the well plate, centrifuged, and used for sacrificial embedded 3D printing. After printing, the constructs were left to gel at 37°C for 20 min. The wells were then filled with cold 100 mM CaCl_2,_ and the plate was left in a 4°C refrigerator for 10 min to allow complete alginate gelation and the Pluronic is liquify. To quantify channel diameters, thin slices (∼1 mm in thickness) of single-channel constructs were taken using a razor blade and imaged under a phase contrast microscope (VWR). Measurements for the area and perimeter of channels were obtained using Fiji software (version 1.54f) which were used to calculate the hydraulic diameter. A relationship, derived from a mass balance around the printing nozzle^81^, was used to relate the hydraulic diameter of printed channels and user-controlled printing parameters (**Equation 2**).

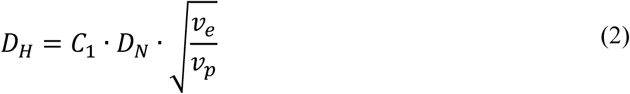

Here, 𝐷_𝐻_ represents the hydraulic diameter of the leftover channel after fully gelling the alginate matrix and evacuating the sacrificial ink, 𝐶_1_ is a constant that captures the combined effects of the printing needle’s inner diameter and alginate swelling on the printed strand dimensions., 𝑣_𝑒_ is the extrusion speed, 𝑣_𝑝_ is the translational speed of the printing needle, and 𝑏 is a correction factor accounting for phenomena such as evaporation, filament contraction, and swelling.

### Print fidelity analysis

Print quality was further assessed by comparing 3D printed channel measurements with the intended geometry. First, alginate partially gelled with 15.0 mM of calcium was prepared as described in section 0, transferred into a flat-bottom 25-mL tube (Sarstedt, 609922115), mixed with 10% v/v HEPES saline (resulting in a 3% w/v final alginate concentration), centrifuged for 2 min at 300 x g, and gently mixed with additional GDL solution, taking care to prevent the formation of air bubbles. The gel was then scooped into a custom tissue holder (2.5 mL for cell line constructs and 1.5 mL for SC-derived islet constructs) and leveled with a metal spatula. A 10-branch vascular template was then 3D printed directly into the alginate with 35 % w/v Pluronic F127. The construct was left to gel at 37°C for 20 min. Afterwards, the alginate and tissue holder were placed into a cold 100 CaCl_2_ bath and left in on ice for 10 min. Constructs were then shipped and stored within 25-mL flat-bottom tubes that were filled with a cold 45 mM CaCl_2_ solution supplemented with 100 ppm sodium azide (MilliporeSigma, S8032) and sealed with Parafilm. The final dimensions of the alginate constructs irrigated with 10-branch perfusable networks were measured by X-ray micro-computed tomography (μCT, eXplore Locus 80; GE Healthcare). Scans were performed at 40 kVp (current: 100 mA) with 100 ms exposure time and 2 × 2 detector binning for a voxel resolution of 45 μm with three frames averaging. The final images were reconstructed using the Parallax Innovations reconstruction tool (Parallax Innovations, Ilderton, ON, Canada). A 3-pixel range gaussian filter was applied to smooth the images and remove artefacts. To obtain the final dimensions of the perfusable network, the negative of the μCT image was taken, followed by a segmentation with 100% quality factor to generate 3D models of the hydrogels. Next, a theoretical 3D model (stl. extension) of the 10 branching channels was created with TinkerCAD (AutoDesk, San Francisco, CA, USA) software and used as the numeric reference of the 3D printed perfusable network. Geometrical conformity assessment was performed using open-source CloudCompare software (version 2.12.4). The 3D reconstruction of the perfusable network and numeric reference 3D model were then superimposed automatically using the “fine cloud registration” tool to quantitatively assess the print fidelity and geometric conformity. Numerical deviations, defined as the closest distance between the points of the 3D image of the hydrogel obtained by segmentation of the uCT image and the reference model, were calculated and assigned to each point). A heat map was generated detailing deviations from -0.4 mm to 0.4 mm.

### Cell line culture

Mouse insulinoma 6 (MIN6) cells (kind gift from Dr. Jun-ichi Miyazaki, Osaka University, received at passage 17, master bank at passage 19, and working banks at passage 27) were cultured in tissue culture treated polystyrene T-flasks (Sarstedt, 833912002) at 37°C and 5% CO_2_ ^82^. The cells were cultured in Dulbecco’s modified Eagle’s medium (Fisher Scientific, 10313021) supplemented with 10% fetal bovine serum (FBS, Fisher Scientific, SH3039603), 100 U/mL penicillin/streptomycin (Fisher Scientific, 15140122), 2 mM L-glutamine (Fisher Scientific, 25030081), and 50 μM β-mercaptoethanol (MilliporeSigma, M6250). Medium was supplied at 0.14 mL/cm^2^ and was changed every 2 days until cells reached ∼85% confluency. Cells were then passaged or harvested using TrypLE Express Enzyme (Fisher Scientific, 12605010). Passage numbers were kept below 40 and we used cell suspensions with a viability of at least 85%, measured using trypan blue staining (MilliporeSigma, T8154) and automated cell counting (TC20, Bio-Rad, 1450102).

The conditionally immortalized βTC-tet cell line which undergoes growth arrest upon activation of the tetracycline operator (ATCC, CRL-3237, received at unknown passage x, master bank at passage x+3, and working banks at passage x+7) was cultured in Dulbecco’s modified Eagle’s medium supplemented with 2.5% FBS, 15% heat-inactivated horse serum (Thermo Fisher Scientific, 26050088), 100 U/mL penicillin/streptomycin, and 2 mM L-glutamine. Medium was supplied at 0.14 mL/cm^2^ and changed every 2 days until cells reached ∼90% confluency^83^. Cells were then passaged or harvested using TrypLE Express Enzyme. Passage numbers were kept below x+20.

The AggreWell400 system (StemCell Technologies, 34425) was used to generate beta cell clusters from cell lines according to manufacturer’s instructions. Cells were seeded at 500 or 1000 cells per microwell, collected 2 days after seeding into the microwells, with non-aggregated cells removed using a 40-µm cell strainer prior to use in experiments.

### Stem cell culture and differentiation

Human embryonic stem cells (hESCs, WA01, H1) were obtained from the WiCell Research Institute (received at passage 17, master bank at passage 19, and working banks at passage 27) with approval from the Canadian Institutes of Health Research’s Stem Cell Oversight Committee_hESCs were cultured in feeder-free conditions on 6-well plates (Sarstedt, 833920) coated with Matrigel (Corning, 354277) in mTeSR medium (Stem Cell Technologies, 85850) and passaged using 50 µM EDTA (0.01 mL/cm^2^, Thermo Fisher Scientific, 15575020) in phosphate-buffered saline (Thermo Fisher Scientific, 14190250) for 8 minutes using a 1:8 to 1:20 split, depending on growth kinetics. Medium was changed daily.

Pancreatic differentiation of H1 ESCs was conducted through a protocol adapted from previous studies ^57,84,85^. Cells were dissociated after reaching 70-80% confluency using TrypLE (0.01 mL/cm^2^, 3 min incubation, Gibco, 12605036) and seeded as single cells at 1.45×105 cells/cm2 on growth factor-reduced Matrigel-coated wells in mTeSR medium supplemented with 10 µM Rho-Associated kinase inhibitor (StemCell Technologies, 72308). **Table S3** outlines the basal medium used in each stage and **Table S4** describes the culturing details and growth factors for each stage while **Table S5** details the supplier information. After stage 4, cells were trypsinized and aggregated (1000 cells/cluster) using AggreWell400 plates. Clusters were transferred to ultralow attachment plates (MilliporeSigma, Corning, CLS3471) at the beginning of S6 and placed on a shaker at 100 rpm. On day 1 of stage 7, clusters were either kept in suspension, used for sacrificial embedded 3D printing or immobilized within 3% w/v alginate and cultured for 25 more days with medium changes occurring every 2 days.

Throughout the differentiation process, cells are diligently monitored for the expression of key markers including SOX17 (stage 1), PDX1 (stage 4), NKX6.1 (stage 4), NeuroD (stage 7), Glucagon (stage 7), and C-peptide (stage 7) using flow cytometry (BD Acuri).

### Perfusable tissue constructs and perfusion culture

The embedded 3D printing matrix was prepared as described in section 0 using 17.5 mM or 15.0 mM of calcium for partial gelation and a cell suspension instead of additional HEPES saline.MIN6 constructs were seeded at 40 × 10⁶ cells/mL to create a high-demand metabolic condition, whereas SC-islet constructs were seeded at ≈ 7.5 × 10⁶ cells/mL. These cell seeding densities were determined based on anticipated therapeutic cell doses, ranges of potential device volumes, and equivalent volumetric oxygen consumption rates. After the cold CaCl_2_ bath, constructs were immediately loaded into a custom flow device (citation) and cultured under perfusion at a flow rate of 3 mL/min/channel in a 37°C and 5% CO_2_ incubator for 2 to 25 days. The medium reservoir (80 mL for cell lines and 50 mL for stem cell-derived cells) was changed every 2 days. Completed medium used for perfusion culture was supplemented with and additional 5 mM of CaCl_2_ to prevent de-gelling. After the culturing period, constructs were subject to glucose stimulated insulin secretion (GSIS) and/or testing end-point viability and cell characterization analysis.

### Live/dead staining

Thin frontal sections (∼1 mm thickness) were manually sliced using a razor blade and then incubated in a HEPES-buffered staining solution containing 5 mM calcium chloride, 10% v/v serum-free cell medium, 4 μM calcein AM (live stain, Thermo Fisher Scientific, C1430), and 30 μM propidium iodide (dead stain, Thermo Fisher Scientific, P1304MP) for 30 min in a 37°C and 5% CO_2_ incubator. The slices were then rinsed with HEPES buffered saline and imaged using an Olympus IX81 inverted microscope system and MetaMorph (version 7.6) acquisition software. Whole slices were imaged using an automated stage system that captured sequential images in a grid pattern. These pictures were then compiled together using ImageJ software (version 1.9.0, Grid/Collection stitching plugin) and processed by subtracting a Gaussian blur-filtered (sigma 200) copy of the compiled image from the original and using the automated Brightness/Contrast adjustment feature.

### Dithizone Staining of Islet Aggregates

Dithizone (DTZ, Sigma, 43820) was prepared fresh before each experiment. A 5 mg/mL stock solution was made by dissolving 5 mg DTZ powder in 1 mL dimethyl sulfoxide (DMSO, Fisher scientific, BP231-100). The stock solution was diluted with 4 mL PBS⁻ (Thermo fisher scientific, 14190250) and mixed thoroughly to dissolve as much dye as possible. The solution was filter-sterilized using a 0.22 µm syringe filter (Fisher scientific, 13100106) and used within 24 h. For staining, a minimum of 10 islet aggregates were collected in a microcentrifuge tube and incubated in 100 µL of the DTZ solution for <1 min. The staining solution was immediately removed, and the aggregates were washed repeatedly with PBS⁻ until the wash solution appeared completely clear. Aggregates were then transferred into a 6-well plate containing 2 mL PBS⁻ for imaging.

### Immunofluorescence Staining of Paraffin-Embedded Tissue Sections

Paraffin-embedded tissue sections were processed for immunofluorescence staining using an in-house 10X PBS buffer (1.37 M NaCl, 27 mM KCl, 100 mM Na₂HPO₄, and 18 mM KH₂PO₄ in 1 L of reverse osmosis (RO) water). For deparaffinization, slides were first incubated at 55 °C for 1 h, then immersed sequentially in xylene (3 × 5 min), 100% ethanol (2 × 10 min), 95% ethanol (2 × 10 min), 70% ethanol (2 × 10 min), and finally rinsed in RO water (2 × 5 min). Antigen retrieval was performed by preparing a 0.5 M Tris buffer (36.34 g Tris base (Fisher scientific, BP152-1) in 600 mL RO water), heating ∼500 mL of the solution to boiling in a microwave during the water rinse step, and immersing the slides in the boiling solution for 17 min, followed by rinsing under tap water for 5 min. Slides were permeabilized in 1X PBS containing 0.1% Triton X-100 (Sigma Aldrich, X100-5ml) and 1% BSA (200 mL PBS, 200 µL Triton, 2 g BSA) for 2 × 10 min. Blocking was performed by encircling tissue sections with a hydrophobic barrier pen (Sigma Aldrich,Z672548-1EA), placing the slides in a humidified chamber, and applying 40 µL of blocking buffer (2 mL 1X PBS, 2 µL Triton X-100, 100 mg BSA, corresponding to 5% BSA) per section for 1 h at room temperature. Primary antibody incubation was carried out without intermediate washing by replacing the blocking buffer with 40 µL of primary antibody solution diluted 1:50–1:200 in antibody diluent (5 mL 1X PBS, 5 µL Triton X-100, 50 mg BSA, corresponding to 1% BSA). Slides were incubated overnight at 4 °C in a humidified chamber. The following day, slides were washed twice for 10 min each in rinsing buffer (400 mL 1X PBS, 400 µL Triton X-100, 400 mg BSA, corresponding to 0.1% BSA). Hydrophobic barriers were reapplied if necessary, and 40 µL of secondary antibody solution (1:400 dilution in antibody diluent) was added to each section. Slides were incubated for 1 h at room temperature in a humidified chamber, followed by two additional 10 min washes in rinsing buffer. Nuclear counterstaining and mounting were performed by adding 3–4 drops of Vectashield mounting medium containing DAPI (Cedarlane (biolynx), VECTH1200), placing coverslips carefully to avoid bubbles, and storing the slides at 4°C until imaging.

### Flow Cytometry Analysis of Differentiation Stages

Cells at Stage 1, Stage 4, and Stage 7 of differentiation were analyzed by flow cytometry to assess lineage-specific marker expression. For all stages, cells were collected in 96-well plates and centrifuged at 300 × g for 5 min, then resuspended in 200 μL/well of FACS buffer (PBS⁻ supplemented with 2% FBS). After a second centrifugation, live/dead staining was performed using the LIVE/DEAD® Fixable Violet Dead Cell Stain Kit (Thermo Fisher). Briefly, 1 μL of a 1:10 diluted dye was added to 100 μL of FACS buffer in each well, excluding unstained controls, and cells were incubated in the dark at room temperature for 30 min. Following incubation, cells were washed once with FACS buffer and fixed in cold Fixation and Permeabilization Solution (BD Cytofix/Cytoperm, BD Biosciences) for 10 min. Cells were then centrifuged at 400 × g for 4 min, the supernatant was discarded, and pellets were resuspended in 200 μL of Perm/Wash buffer (BD Biosciences).

Intracellular staining was performed using antibodies corresponding to differentiation stage. For Stage 1 definitive endoderm, cells were stained for SOX17 (PE Mouse anti-Human Sox17, BD, Cat. No. 561591, 1:333 dilution). For Stage 4 pancreatic progenitors, cells were stained with NKX6.1 (Alexa Fluor® 647 mouse anti-NKX6.1, BD, Cat. No. 563338, 1:33 dilution) and PDX1 (PE mouse anti-PDX1, BD, Cat. No. 562161, 1:33 dilution), with matched isotype controls. For Stage 7 endocrine cells, cells were stained with antibodies against glucagon (PE Mouse Anti-Glucagon, BD, Cat. No. 565860, 1:3333 dilution) and C-peptide (AF647 mouse anti-C-peptide, BD, Cat. No. 565831, 1:3333 dilution), alongside isotype controls. After antibody incubation, cells were washed in Perm/Wash buffer, resuspended in FACS buffer, and stored protected from light on ice or at 4 °C until acquisition. Flow cytometry was performed on a BD Accuri C6 cytometer, and data were analyzed using standard gating strategies to assess marker expression and live/dead exclusion.

### Glucose stimulated insulin secretion

For static assays, cell clusters and fragments of tissue constructs (∼200 μL) were incubated in wells of a 12-well plate containing Kreb’s buffer (2 mL/well) with different glucose (MilliporeSigma, G8270) and potassium chloride (KCl, MilliporeSigma, P5405) concentrations (Supplementary **Table S6**). Kreb’s buffer (pH 7.4) was prepared with 130 mM sodium chloride (MilliporeSigma, S5886), 4.7 mM KCl, 1.2 mM magnesium sulfate (MgSO_4_, MilliporeSigma, M2643), 10 mM HEPES, 2.5 mM calcium chloride dihydrate (CaCl_2_•2H_2_O, MilliporeSigma, C3306), 5.0 mM sodium bicarbonate (NaHCO_3_, MilliporeSigma, S5761) and 0.5% w/v bovine serum albumin (BSA, MilliporeSigma, A7906) in reverse osmosis water. First, solutions were warmed to 37°C within a cell culture incubator. Next, cell clusters and tissue fragments were transferred between wells within a 40-μm cell strainer and incubated according to **Table S6**. Solution samples after each stage were centrifuged at 300 x g for 5 min to remove possible debris and the supernatant was then collected and stored frozen at -20°C until analysis.

To test the dynamic GSIS function of beta cells within perfusable alginate constructs, the tubing of the perfusion system was rearranged into a single-pass configuration where the liquid leaving the flow device would be captured into sampling tubes instead of returning to the reservoir and cycled around. Next, the loop was flushed of cell culture medium using low-glucose Kreb’s buffer. Next, the dynamic GSIS profiles of constructs were determined using five stages by exchanging the perfusion system reservoir: 1) a synchronization stage in low-glucose Kreb’s; 2) a basal stage in low-glucose Kreb’s; 3) a stimulation stage in high glucose Kreb’s; 4) a second basal stage at low glucose; 5) a decoupling stage in KCl Kreb’s buffer. During stages 2-5, the perfusate at the outlet was continuously collected in 50-mL tubes with sampling tubes being rotated out every 5 min for cell line constructs and 3 min for SC-derived islet constructs. The tubes were then centrifuged at 300 x g for 5 min and a 500-μL sample of the supernatant was then collected and stored at -20°C until analysis. **Table S6** summarizes the duration and perfusate details of each stage. The insulin measurements for cell line experiments were quantified using a mouse insulin ELISA kit (Cedarlane, Alpco, 80-INSMS-E01) while the C-peptide in samples from stem cell experiments were measured using a human C-peptide ELISA kit (Cedarlane, Alpco, 80-CPTHU-E01.1) according to the manufacturer’s instructions. The optical absorbance for these assays was measured using a Benchmark Plus Microplate Spectrophotometer (Bio-Rad). For static assays, the stimulation index was calculated by dividing the C-peptide reading from the stimulation condition by that of the first basal condition.

### Oxygen transport modelling

Theoretical oxygen profiles were generated using COMSOL Multiphysics 5.5 software as previously described ^86^. Here, we simplify the model by excluding fluid dynamics. Briefly, only radial oxygen transport, a constant oxygen tension (140 mmHg) at the channel walls, and no diffusion at the outer edges of the tissue slice were assumed. Oxygen diffusivity was estimated as a weighted average of the diffusivity of oxygen through alginate gel and cells and Monod oxygen consumption kinetics were adopted. The coordinates of channels for multi-branched were modelled after microscope images and channel cross-sections were approximated as circles. **Table S7** summarizes the model parameters.

### Statistical Analysis

GraphPad Prism software (version 10.2.2) was used to conduct the statistical analysis and for model fitting. First, data sets were examined for normality using the Shapiro-Wilk test. One-way ANOVA with Tukey post-hoc testing with a threshold of p < 0.05 was used to determine partial gelation effects while a series of unpaired t-tests were conducted to compare different conditions. Practical equivalency in mechanical properties and viability was determined using two one-sided t tests with a threshold of p < 0.05 for differences of >5% of the means. The standard error of the mean (SEM) for line plots are presented as shaded areas. Reported values and bar graph data represent mean ± SEM. All experiments were conducted in triplicate (n=3) unless otherwise stated. Statistical significance is reported using the following notation: *P<0.05, **P<0.01, and ***P<0.001.

## Acknowledgements

We would like to thank the lab of Prof. Matt Kinsella at McGill University and Dr. Maedeh Rahimnejad who recently graduated from Université de Montréal for their collaboration and guidance on this project. This research was made possible by the funding support provided by the Fonds de recherche du Québec – nature et technologies (FRQ-NT 2022-PR-297722), the New Frontiers in Research Fund (NFRFE-2018-01546), Médicament Québec, Natural Sciences and Engineering Research Council of Canada (NSERC), the Canadian Foundation for Innovation (35507) and the Canada Research Chairs program (C.H.). We also acknowledge the support provided by the following networks: the Quebec Cell, Tissue and Gene Therapy Network–ThéCell and the Cardiometabolic Health, Diabetes and Obesity – CMDO Research Network (thematic networks supported by the Fonds de recherche du Québec–Santé), PROTEO (The Quebec Network for Research on Protein Function), and CQMF/QCAM (Quebec Centre for Advanced Materials).

## Ethics declarations

C.H. is Chief Scientific Officer and co-founder of CellTerix Biomedical, a company developing devices for diabetes therapy. J.B. is Chief Executive Officer and co-founder of CellTerix Biomedical.

## Contributions

BNM contributed to conceptualization, data curation, formal analysis, investigation, investigation, methodology, project administration, resources, software, validation, visualization, writing – original draft, and writing – review & editing.

HEO contributed to data curation, formal analysis, investigation, methodology, project administration, resources, software, validation, visualization, writing – original draft, and writing – review & editing.

TL contributed to data curation, formal analysis, investigation, methodology, validation, visualization, writing – original draft, and writing – review & editing.

JB contributed to methodology, resources, validation, and writing – review & editing.

SP contributed to funding acquisition, resources, supervision, and writing – review & editing. SL contributed to funding acquisition, resources, supervision, and writing – review & editing. MAF contributed to funding acquisition, resources, supervision, and writing – review & editing.

RLL contributed to conceptualization, funding acquisition, resources, supervision, and writing – review & editing.

CAH contributed to conceptualization, data curation, funding acquisition, methodology, project administration, resources, supervision, and writing – review & editing.

---

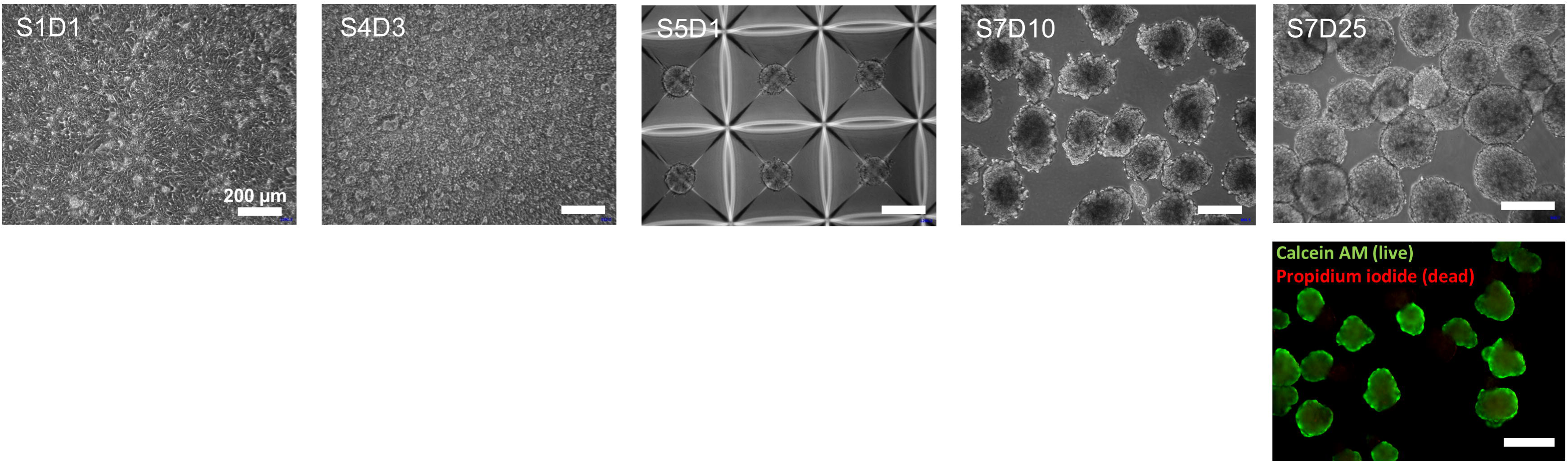

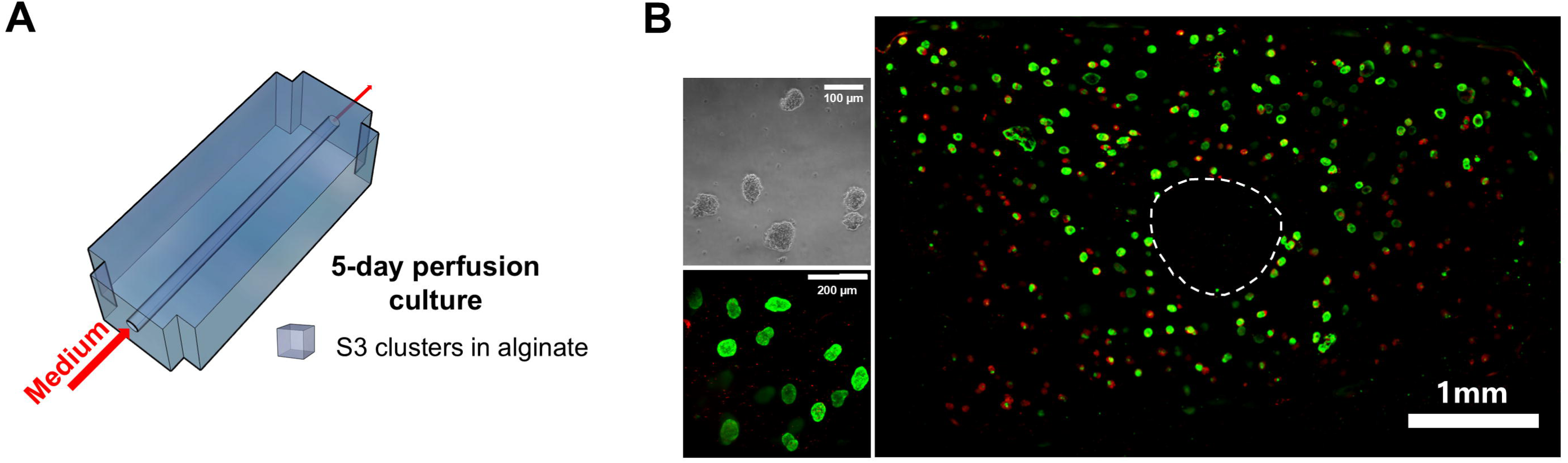

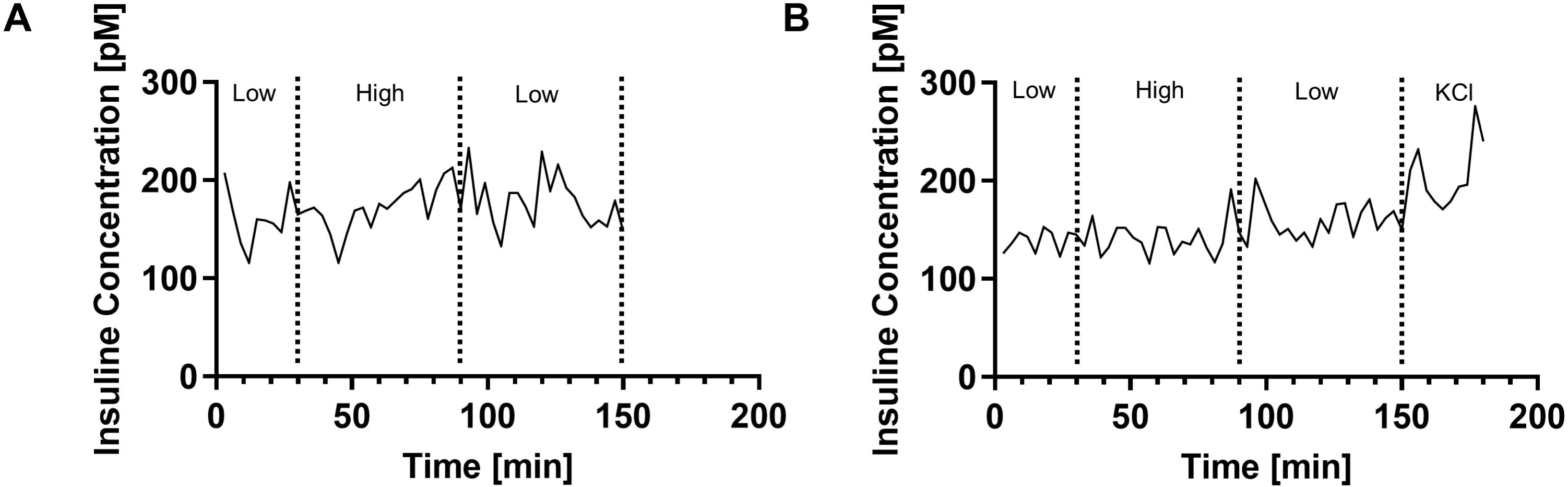

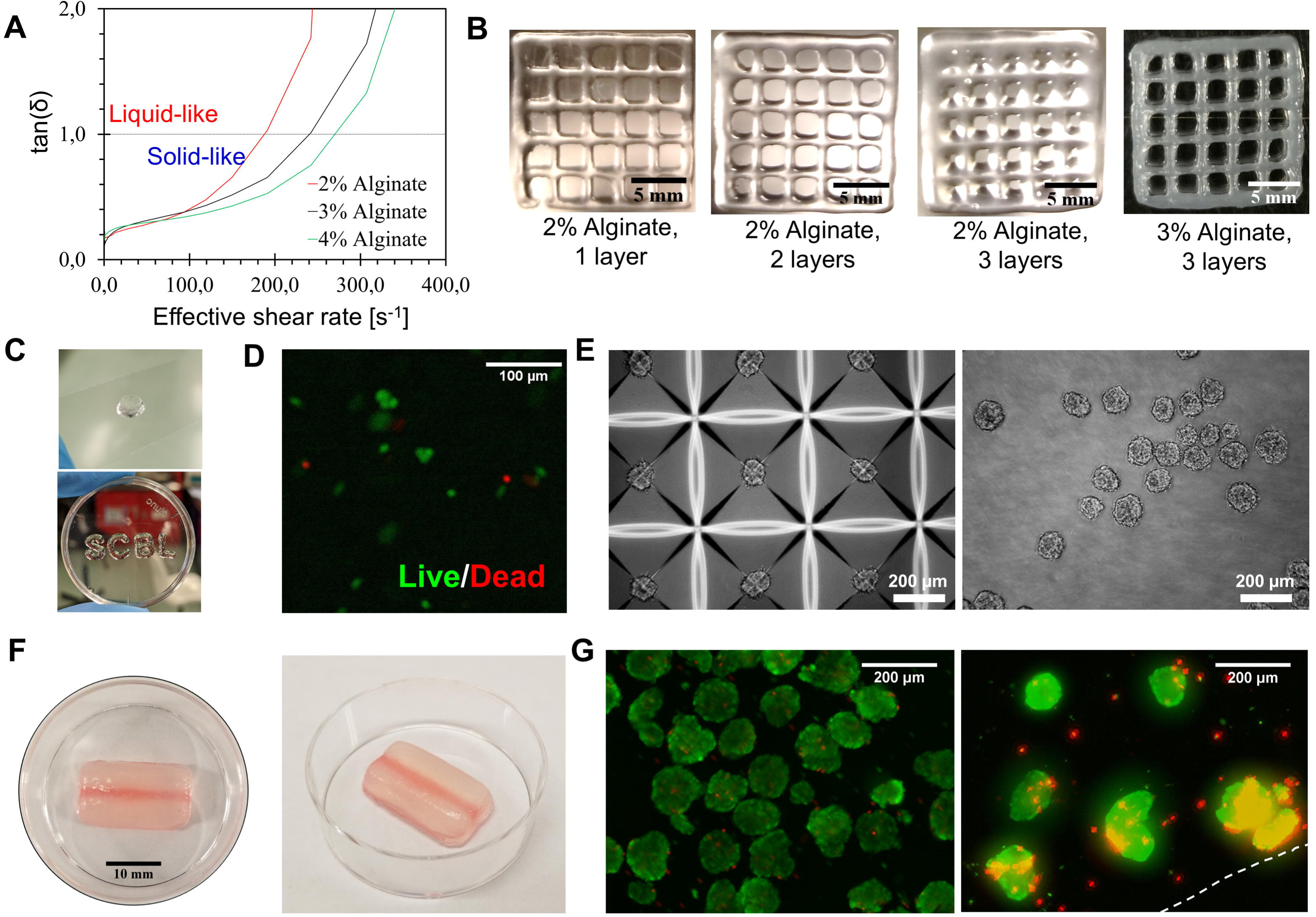

