## Supplementary for "Extended perfused culture of cm-scale endocrine pancreatic tissues created through sacrificial embedded printing into alginate"

**Table S1.** Herschel Bulkley parameters of partially gelled alginate

| Calcium concentration [mM] | Yield stress, $\tau_0$ [Pa] | Flow coefficient, k [Pa·s] | Herschel-Bulkley index, n [-] |
|---|---|---|---|
| 10.0 | 11.0 ± 0.6 | 12.9 ± 0.5 | 0.48 ± 0.01 |
| 12.5 | 22.9 ± 0.8 | 19.8 ± 0.7 | 0.44 ± 0.01 |
| 15.0 | 36 ± 1 | 15.6 ± 0.8 | 0.50 ± 0.01 |
| 17.5 | 61.9 ± 0.5 | 13.0 ± 0.3 | 0.54 ± 0.01 |
| 20.0 | 61 ± 2 | 14 ± 2 | 0.53 ± 0.02 |
| 25.0 | 51 ± 4 | 18 ± 3 | 0.46 ± 0.03 |

**Table S2**: Comparison of required islet and MIN6-equivalent cell densities for a 720,000-IEQ dose distributed in device volumes of 4 mL and 2 mL under two metabolic demand assumptions.

| Model | Weight (kg) | IEQ/kg | Total IEQ | Device Volume (mL) | IEQ/mL | Islet Cells (M/mL) | MIN6 Eq. (M/mL) | Islet Cells Final (M/mL) | MIN6 Final (M/mL) |
|---|---|---|---|---|---|---|---|---|---|
| **Moderate metabolic demand (50%)** | 90 | 8000 | 720,000 | 4 (10 channel device) | 180,000 | 317.7 | 83.7 | 158.85 | 41.85 |
| **Low metabolic demand (1%)** | 90 | 8000 | 720,000 | 2 (5-channel device) | 360,000 | 635.4 | 167.5 | 6.354 | 1.675 |

**Table S3.** Basal medium for pancreatic differentiation of ESCs

| Stage | Basal Medium |
|---|---|
| 0 | mTeSR1 |
| 1-2 | MCDB131 + 10 mM D-glucose + 1.5 g/L sodium bicarbonate (NaHCO3) + 0.5% fatty acid-free bovine serum albumin (FAF-BSA) + 1X GlutaMAX supplement |
| 3-4 | MCDB131 + 15 mM D-glucose + 2.56 g/L NaHCO3 + 2% FAF-BSA + 1X GlutaMAX supplement + 1X Insulin-transferrin-selenium-ethanolamine (ITS-X) supplement + 250 µM Ascorbic Acid (AA) |
| 5-6 | MCDB131 + 20 mM D-glucose + 1.5 g/L NaHCO3 + 2% FAF-BSA + 1X GlutaMAX supplement + 1X ITS-X supplement + 10 µg/mL heparin + 10 µM zinc sulfate + 100 U/mL pen/strep |
| 7 | MCDB131 + 1.5 g/L NaHCO3 + 2% FAF-BSA + 1X GlutaMAX supplement + 1X ITS-X supplement + 10 µg/mL heparin + 10 µM zinc sulfate + 100 U/mL pen/strep |

**Table S4.** Static pancreatic differentiation of ESCs parameters

| Stage | | Time [days] | Culture details | | | Additives |
|---|---|---|---|---|---|---|
| | | | Culture format | Medium volume ratio | Medium changes [days] | |
| - | 0 | ~2 | Matrigel-coated plates | 0.2 mL/cm2 | 1 | None |
| hESCs | 1A | 1 | Matrigel-coated plates | 0.2 mL/cm2 | 1 | 3 µM CHIR 99021<br>100 ng/mL Act A |
| | 1B | 2 | Matrigel-coated plates | 0.2 mL/cm2 | 1 | 100 ng/mL Act A |

| | | | | | | |
|---|---|---|---|---|---|---|
| Definitive Endoderm (DE) | 2 | 3 | Matrigel-coated plates | 0.2 mL/cm2 | 1 | 50 ng/mL KGF<br>1250 nM IWP-2<br>250μM AA |
| Primitive gut tube (PGT) | 3 | 2 | Matrigel-coated plates | 0.2 mL/cm2 | 1 | 50 ng/mL KGF<br>250 nM SANT-1<br>100 nM LDN<br>200 nM TPB<br>1 μM RA |
| Pancreatic endoderm (PE) | 4A | 3 | Matrigel-coated plates | 0.2 mL/cm2 | 1 | 50 ng/mL KGF<br>250 nM SANT-1<br>200 nM LDN<br>100 nM TPB<br>0.1 μM RA |
| | 4B | 1 | AggreWell plate | 5 mL/well | 1 | 50 ng/mL KGF<br>250 nM SANT-1<br>200 nM LDN<br>100 nM TPB<br>0.1 μM RA |
| Endocrine precursors (EP) | 5 | 4 | AggreWell plate | 5 mL/well | 1 | 250 nM SANT-1<br>100 nM γSiXX<br>1 μM T3<br>10 μM Alk5ii<br>μM RA<br>100 nM LDN |
| Immature beta cells (iBC) | 6 | 7 | 6-well plate, 100 RPM | 5 mL/well | 1 | 100 nM γSiXX<br>μM T3<br>10 μM Alk5ii<br>100 nM LDN |

| | | | | | | |
|---|---|---|---|---|---|---|
| Mature beta cells (mBC) | 7 | 25 | 6-well plate, 100 RPM | 5 mL/well | 2 | 1 μM T3 |

Act A: activin A; KGF: Recombinant Human KGF/FGF-7 Protein; AA: L-ascorbic acid; LDN: LDN193189; TPB: TPB (PKC activator); RA: retinoic acid; γSiXX: γ-secretase inhibitor XX; T3: T3 (3,3′,5-Triiodo-L-thyronine sodium salt); Alk5ii: ALK5 Inhibitor II.

**Table S5**: Supplier information for pancreatic differentiation reagents

| Reagent | Supplier | Catalog number |
|---|---|---|
| MCDB 131 | Thermo Fisher Scientific | 10372019 |
| D-glucose | MilliporeSigma | G8769 |
| NaHCO3 | Fisher | S233500 |
| FAF-BSA | Proliant | 68700 |
| GlutaMAX™ supplement | Thermo Fisher Scientific | 35050061 |
| AA | MilliporeSigma | A4544 |
| ITS-X supplement | Thermo Fisher Scientific | 51500056 |
| heparin | MilliporeSigma | H3149100ku |
| zinc sulfate | Fisher | 7733020 |
| CHIR 99021 | Cedarlane | 4423/10 |
| Activin A | Cedarlane | 338AC50/CF |
| KGF | StemCell Technologies | 781862 |
| IWP-2 | Cedarlane | 13951 |
| SANT-1 | MilliporeSigma | S4572 |
| LDN | MilliporeSigma | SML0559 |
| TPB | MilliporeSigma | 565740 |
| RA | MilliporeSigma | R2625 |
| γSiXX | MilliporeSigma | A4544 |
| T3 | MilliporeSigma | T6397 |
| Alk5ii | Cedarlane | 14794 |

**Table S6.** Glucose stimulated insulin secretion assay parameters

| Stage | Additions to Kreb's buffer | Static incubation time [min] | Dynamic perfusion time [min] |
|---|---|---|---|
| Synchronization | 2.8 mM glucose | 60 | 120 |
| First low glucose | 2.8 mM glucose | 60 | 30 |
| High glucose | 16.7 mM glucose | 60 | 60 |
| Second low glucose | 2.8 mM glucose | 60 | 60 |
| Depolarization | 20 mM KCl | 30 | 30 |
| Collection frequency | N/A | N/A | 5 |

**Table S7.** Oxygen transport parameters for computational modelling

| Parameters | Value | Source |
|---|---|---|
| Maximum oxygen consumption rate, MIN6 ($R_{O_2,M6}$) | 0.129 mol s-1 m-3 | 87–90 |
| Maximum oxygen consumption rate, βTC-tet ($R_{O_2,TC}$) | 0.064 mol s-1 m-3 | 91,92 |
| Necrotic oxygen tension ($C_{cr}$) | 0.1 mmHg | 93–95 |
| Monod constant for oxygen consumption, MIN6 ($K_{s,M6}$) | 0.62 mM | 88,93,96 |
| Monod constant for oxygen consumption, MIN6 ($K_{s,TC}$) | 0.01 mM | 91,92 |
| Oxygen diffusivity in alginate ($D_{alg}$) | 2.54·10-5 cm2 s-1 | 97,98 |
| Oxygen diffusivity in MIN6 cells ($D_{M6}$) | 1.24·10-5 cm2 s-1 | 88,93 |
| Oxygen diffusivity in bTC-tet cells ($D_{TC}$) | 1.62·10-5 cm2 s-1 | 91 |
| Atmospheric oxygen tension | 140 mmHg | 74 |
| Minimum viable oxygen tension | 1.16 mmHg | 9 |

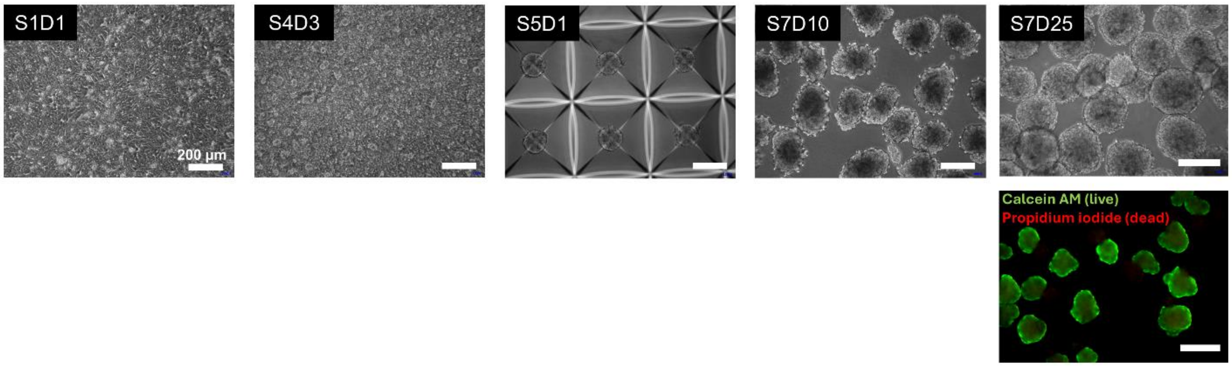


**Figure S1.** Morphological progression and viability of stem cell-derived clusters during differentiation and suspension culture. Phase-contrast images show morphological changes in stem cell-derived clusters at key stages of differentiation under suspension conditions: (S1D1) early definitive endoderm stage, (S4D3) pancreatic progenitor stage, and (S5D1) cell aggregation within AggreWell plates. Continued maturation is observed at (S7D10) and (S7D25), corresponding to day 10 and day 25 of Stage 7 (endocrine differentiation). A live/dead assay using Calcein AM (green, live cells) and Propidium Iodide (red, dead cells) indicates high viability of clusters after 25 days of suspension culture.

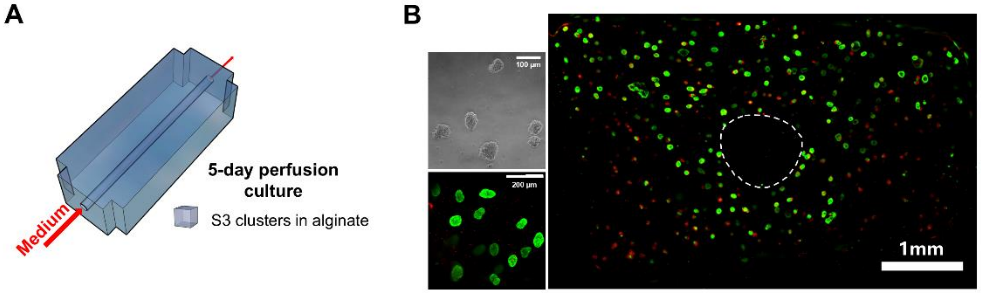


**Figure S2.** Viability of stem cell-derived clusters following short-term perfusion culture in alginate. **(A)** Schematic of the embedded 3D construct containing Stage 3 (S3) cell clusters encapsulated in alginate and cultured under perfusion for 5 days. **(B)** Bright-field and fluorescence images showing the morphology and viability of encapsulated clusters after perfusion. Live/dead staining with Calcein AM (green, live cells) and Propidium Iodide (red, dead cells) reveals high viability across the construct cross-section, with the region corresponding to a printed vascular channel outlined (dashed line).

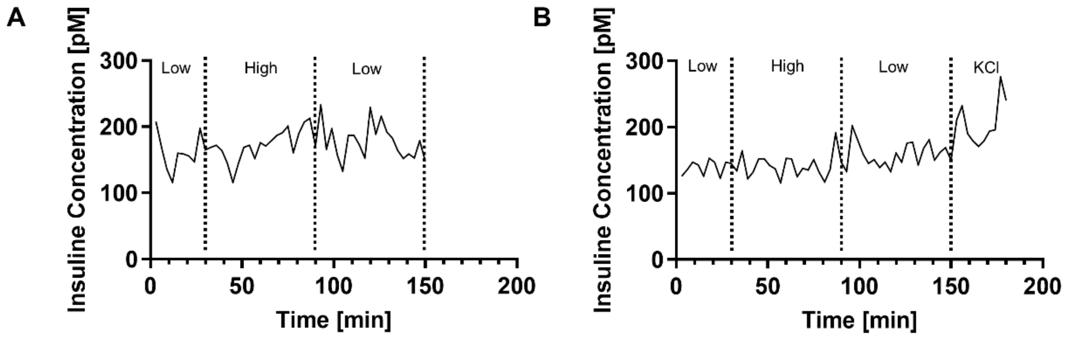


**Figure S3.** Glucose-stimulated insulin secretion (GSIS) response of human islets cultured under perfusion. **(A)** GSIS profile after 2 days of perfusion culture shows an increase in insulin secretion toward the end of the high-glucose stimulation phase. **(B)** The same construct after 7 days of perfusion exhibits a similar response pattern, with insulin release predominantly observed at the end of the high-glucose cycle and a further increase upon KCl stimulation

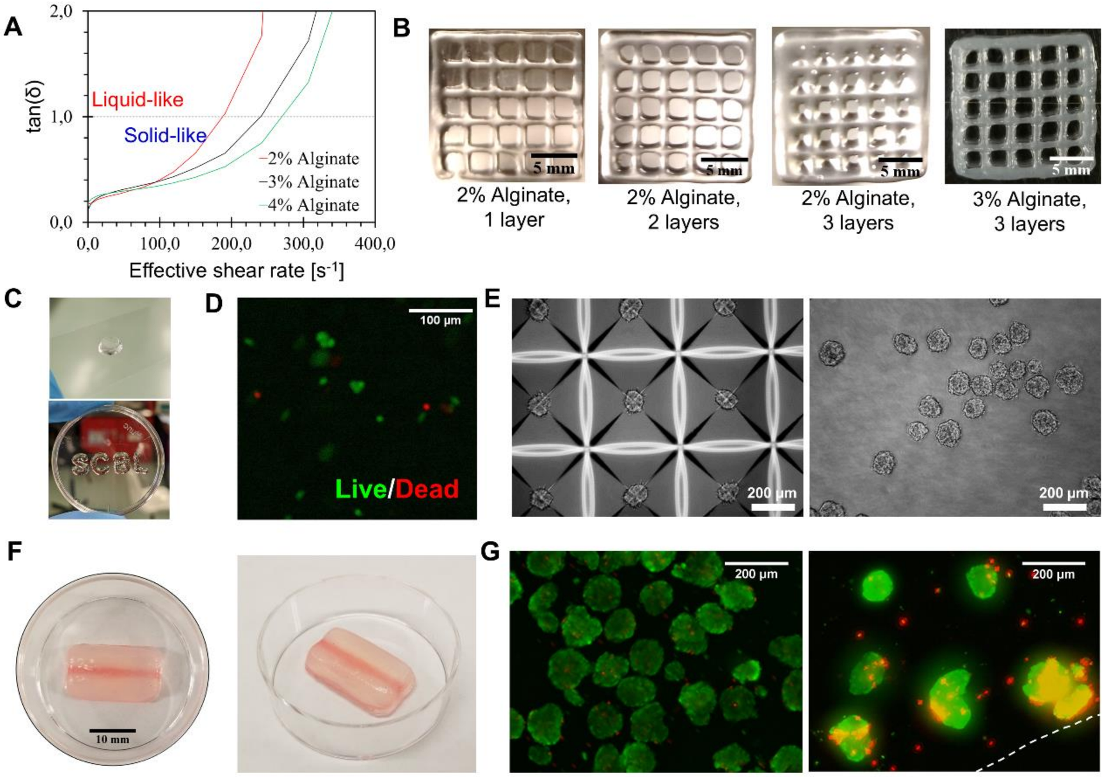


**Figure S4.** Layer-by-layer 3D printing with partially gelled alginate. **(A)** Deflection angle (loss modulus divided by solid modulus) of alginate of different concentrations partially gelled with 17.5 mM of calcium. **(B)** Grid printed using 2% and 3% alginate partially gelled with 17.5 mM calcium. **(C)** Top: cell-laden disk construct 3D printed layer-by-layer with 3% alginate partially gelled with 17.5 mM calcium ($2 \cdot 10^6$ cells/mL). Bottom: acellular letters 3D printed using same ink. **(D)** βTC-tet cell viability in disk construct shown in (C). Live cells are stained green (calcein-AM) and dead cells are stained red (ethidium homodimer-1). **(E)** Left: βTC-tet clusters in Aggrewell plate (250 cells/cluster). Right: suspended βTC-tet clusters. **(F)** Acellular single-channel construct 3D printed layer-by-layer using 3% alginate partially gelled with 17.5 mM Calcium and 38% w/v Pluronic F127. **(G)** Left: βTC-tet cluster viability before bioprinting. Right: βTC-tet cluster viability after layer-by-layer 3D printing in 3% alginate partially gelled with 17.5 mM calcium. The dotted line indicates the edge of the construct.
